# Xmas ESC: A new female embryonic stem cell system that reveals the BAF complex as a key regulator of the establishment of X chromosome inactivation

**DOI:** 10.1101/768507

**Authors:** Andrew Keniry, Natasha Jansz, Linden J. Gearing, Iromi Wanigasuriya, Joseph Chen, Christian M. Nefzger, Peter F. Hickey, Quentin Gouil, Joy Liu, Kelsey A. Breslin, Megan Iminitoff, Tamara Beck, Andres Tapia del Fierro, Lachlan Whitehead, Sarah A. Kinkel, Phillippa C. Taberlay, Tracy Willson, Miha Pakusch, Matthew E. Ritchie, Douglas J. Hilton, Jose M. Polo, Marnie E. Blewitt

## Abstract

Although female pluripotency significantly differs to male, complications with *in vitro* culture of female embryonic stem cells (ESC) have severely limited the use and study of these cells. We report a replenishable female ESC system, Xmas, that has enabled us to optimise a protocol for preserving the XX karyotype. Our protocol also improves male ESC fitness. We utilised our Xmas ESC system to screen for regulators of the female-specific process of X chromosome inactivation, revealing chromatin remodellers Smarcc1 and Smarca4 as key regulators of establishment of X inactivation. The remodellers create a nucleosome depleted region at gene promotors on the inactive X during exit from pluripotency, without which gene silencing fails. Our female ESC system provides a tractable model for XX ESC culture that will expedite study of female pluripotency and has enabled us to discover new features of the female-specific process of X inactivation.

## Introduction

Female pluripotent stem cells differ from males genetically, epigenetically and functionally (Choi et al., 2017a; Choi et al., 2017b; Ooi et al., 2010; Schulz et al., 2014; Yagi et al., 2017; Zvetkova et al., 2005). Despite this, the vast majority of ESC research has been performed on male lines, leading to a substantial imbalance in our understanding of sex-specific pluripotency. The first confirmed ESC line to be derived was male (Bradley et al., 1984). Subsequently, the ESC lines employed as workhorses cells for the field, E14, R1, J1 and Bruce4, were all male (Hooper et al., 1987; Kontgen et al., 1993; Li et al., 1992; Nagy et al., 1993). Strikingly, all of the 13 karyotyped ESC lines commercially available via the American Type Culture Collection (ATCC) are male. This major imbalance has substantially hindered an understanding of how female and male pluripotency may differ and impeded study of female-specific processes in their native context, including X chromosome inactivation.

When cultured in the primed state, female ESCs are epigenetically unstable, displaying global hypomethylation compared to males (Zvetkova et al., 2005). Further widespread loss of DNA methylation, including at repetitive elements and imprinting control centres is observed when female ESCs are cultured in conditions that promote ground-state pluripotency, despite male ESCs tolerating such conditions (Choi et al., 2017a; Choi et al., 2017b; Habibi et al., 2013; Ooi et al., 2010; Schulz et al., 2014; Yagi et al., 2017). Female ESCs are also karyotypically unstable, with XO cells rapidly dominating cultures (Choi et al., 2017b; Yagi et al., 2017; Zvetkova et al., 2005), which is also observed in human female embryonic stem cells (Di Stefano et al., 2018). These complications additionally arise when somatic female cells are reprogrammed to induced pluripotent stem cells (iPSCs) (Milagre et al., 2017; Pasque et al., 2018; Song et al., 2019), suggesting that these are features of female pluripotency in culture that confound both research requiring such cells and future medical applications.

Another distinguishing feature of female ESCs is their unique X chromosome inactivation (XCI) status. XCI is the mammalian compensation mechanism that ensures equal gene dosage between XX females and XY males, resulting in the near complete silencing of one female X chromosome, reviewed in (Brockdorff and Turner, 2015; Disteche and Berletch, 2015; Gendrel and Heard, 2011; Jegu et al., 2017). Female ESCs, like the embryonic cells of the blastocyst from which they are derived, have activity from both X chromosomes. This exclusively occurs in ESCs and primordial germ cells (Kratzer and Chapman, 1981; Monk and McLaren, 1981; Tam et al., 1994). Upon differentiation they undergo random XCI leaving the cell with an active (Xa) and an inactive (Xi) X chromosome where the choice of which parental chromosome becomes the Xi appears random. XCI occurs in a stepwise process after being initiated by the upregulation of the long non-coding RNA *Xist*, which spreads *in cis* to coat the future Xi (Brockdorff et al., 1991; Brown et al., 1992). *Xist* then recruits silencing factors (Chu et al., 2015; McHugh et al., 2015; Minajigi et al., 2015) that establish the Xi through the loss of activating (Heard et al., 2001; Keohane et al., 1996; McHugh et al., 2015; Zylicz et al., 2019) and gain of repressive histone marks (Boggs et al., 2002; de Napoles et al., 2004; Fang et al., 2004; Heard et al., 2001; Keniry et al., 2016; Mak et al., 2002; Mermoud et al., 2002; Peters et al., 2002; Plath et al., 2003; Plath et al., 2004; Schoeftner et al., 2006; Silva et al., 2003) and the adoption of a unique bipartite chromosome confirmation (Deng et al., 2015; Giorgetti et al., 2016; Nora et al., 2012; Splinter et al., 2011). Silencing of the Xi is then maintained by DNA methylation (Keohane et al., 1996; Sado et al., 2000), H3K9me3 (Keniry et al., 2016; Minkovsky et al., 2014) and the chromatin regulator Smchd1 (Blewitt et al., 2008; Gendrel et al., 2012). This rich understanding of the ontogeny of XCI is the result of three decades of exceptional research. However, we still do not completely understand the process of XCI, in part because the major issues with female ESC culture have meant there has not been a tractable system in which to study the early stages on XCI in its native context.

After establishment of XCI, hypomethylation and the XO karyotype are no longer a feature of female cells, with XCI seemingly having a stabilising effect (Schulz, 2017). There is increasing evidence that two active X chromosomes cause female ESCs to behave differently to males, as both female ESCs and iPSCs that have spontaneously become XO have similar transcriptomes, epigenomes and differentiation potential to XY ESCs (Choi et al., 2017b; Pasque et al., 2018; Schulz et al., 2014; Song et al., 2019; Zvetkova et al., 2005). Recently, there has been interest in correcting the stability of female ESCs, resulting in discovery of chemical inhibitors able to preserve the methylome, but not the XX karyotype (Choi et al., 2017b; Yagi et al., 2017). Based on the uniqueness of female pluripotency, the challenges of working with female ESCs, and a desire to study XCI in normal female ESC, we created X-linked reporter alleles (Xmas). The Xmas ESC system allowed us to rapidly monitor both karyotype and XCI status to improve culture conditions and deepen our understanding of female ESC biology. Using the Xmas ESC system, we performed the first screen for regulators of the establishment of XCI in its native state. Our screen revealed a key role for the nucleosome remodellers Smarcc1 and Smarca4 in the establishment of XCI. Smarcc1 creates an accessible future Xi that allows XCI to proceed. This is the first report of chromatin relaxation being an initial step in gene silencing, which shows that by screening in normal female ESC we can reveal new aspects of XCI.

## Results

### Establishment of Xmas ESCs and an improved culture system for female pluripotent stem cells

The rapid outgrowth of XO ESCs in culture presents a challenge to the study of female pluripotency and requires that a replenishable source of XX cells with an easily monitorable karyotype be available (Choi et al., 2017b). To this end, we created two reporter alleles by knock-in of either a GFP or mCherry reporter into the 3’UTR of the X-linked and constitutively expressed house-keeping gene *Hypoxanthine guanine phosphoribosyltransferase* (*Hprt)*, in C57BL/6 ESC (X^*Hprt*-GFP^ and X^*Hprt*-mCherry^, Figure 1A). To ensure we could constantly rederive XX ESCs, we created two homozygous/hemizygous mouse strains from the reporter alleles which when crossed produce female offspring that express GFP and mCherry from different X chromosomes (X^*Hprt*-GFP^ X^*Hprt*-mCherry^) (Figure 1B). We call this the Xmas (X-linked markers active silent) system. Importantly, we inserted an internal ribosome entry site (*IRES*) between the stop codon of *Hprt* and the fluorescent reporters. We also deleted the neomycin selection cassette using FlpE recombinase. These two steps ensured that *Hprt* was transcribed efficiently and the protein was unmodified. We observed roughly equal numbers of male and female pups born from each genotype, suggesting wild-type Hprt function was retained (Figure S1A–D). We first tested that the reporter alleles behaved as expected according to random XCI. Flow cytometry of white blood cells from X^*Hprt*-GFP^ X and X^*Hprt*-mCherry^ X animals showed that approximately half the cells were positive for each fluorescent protein (Figure 1C,D). Secondly, we used flow cytometry to detect the reporter alleles in female X^*Hprt*-GFP^ X^*Hprt*-mCherry^ embryos, produced by intercrossing the two strains. Roughly half the cells were positive for each fluorescent marker, in both *ex vivo* hematopoietic stem and progenitor cells (Figure 1E) and primary mouse embryonic fibroblasts (MEFs) derived from the embryos (Figure 1F). These data suggest the reporter constructs are not influencing random XCI, but rather accurately reflect the XCI state.

**Figure 1.**
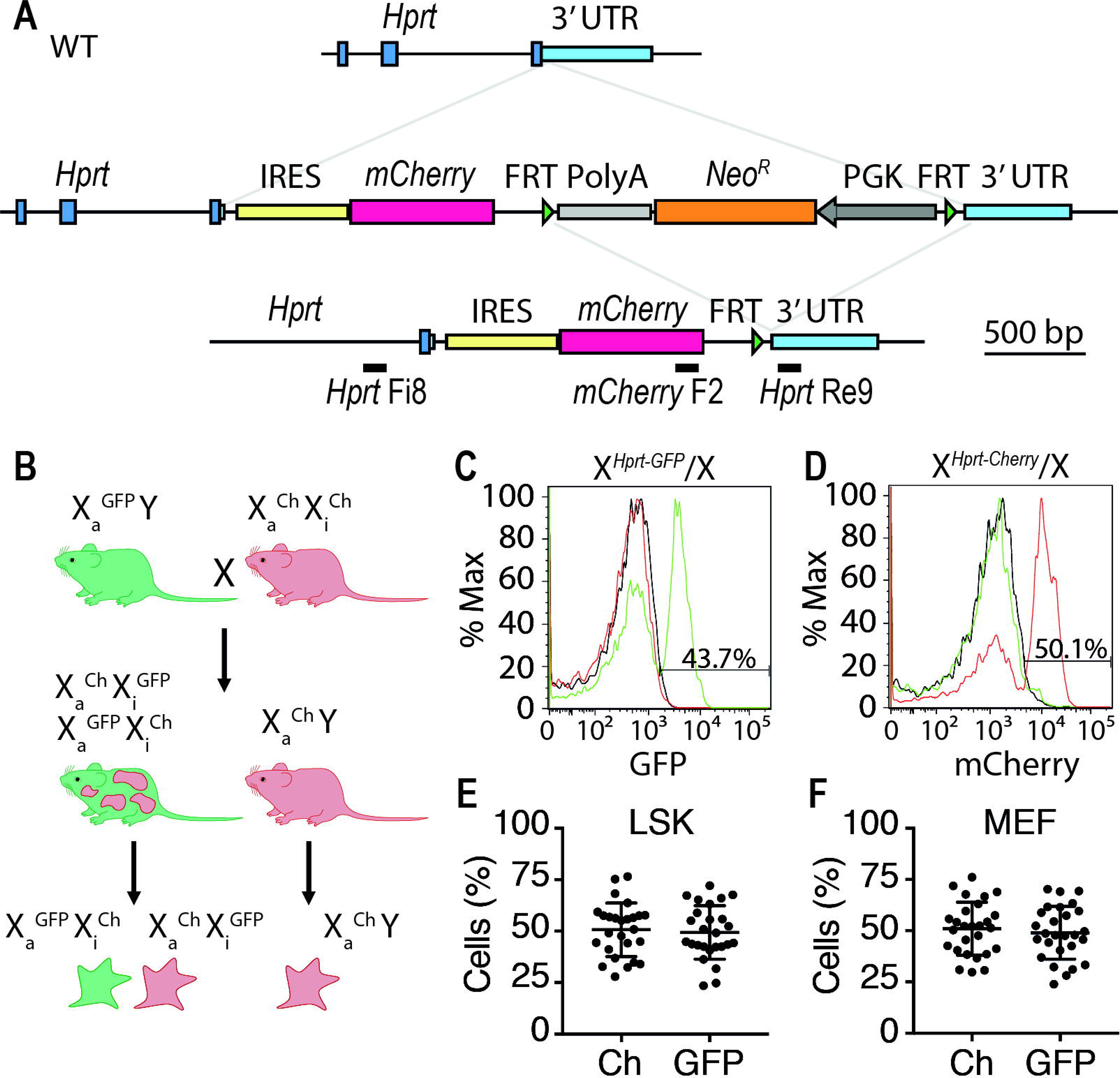
Creation of Xmas reporter alleles and strains of mice. **(A)** Schematic of the mCherry Xmas reporter allele indicating the knockin of the reporter into the 3’ untranslated region (UTR) of *Hprt*. The GFP reporter allele was designed and cloned similarly. The Flipase recognition target (FRT) and internal ribosome entry sites (IRES) are indicated, as are the genotyping oligonucleotides. **(B)** Schematic of homozygous/hemizygous reporter allele mouse strains and their XCI status. (**C,D)** Flow cytometry data showing detection of the GFP **(C)** and mCherry **(D)** fluorescent reporters from white blood cells of X^*Hprt-*GFP^/X (green) and X^*Hprt*-mCherry^/X (red) female mice, compared to XY (black) male mice. **(E,F)** Flow cytometry data showing the percentage of each fluorescent reporter allele from *ex vivo* haematapoetic stem and progenitor cells (LSK) n = 26 **(E)** and primary MEFs n = 26 **(F)**.

We next assessed the suitability of our homozygous mouse lines for production of X^*Hprt*-GFP^ X^*Hprt*-mCherry^ ESCs, Xmas ESCs (Figure 2A). Female blastocysts at embryonic day 3 (E3.5), displayed reporter expression exclusively from the maternal X chromosome in extraembryonic cells, as expected based on the imprinted XCI found in these tissues and in the pre-implantation mouse embryo (Takagi and Sasaki, 1975). By contrast, both X chromosomes were active in the inner cell mass, indicating that reactivation of the silent paternal X chromosome in embryonic tissue occurred as expected (Figure 2B). Following derivation, expression of both reporter alleles was detectable in Xmas ESCs, both by microscopy (Figure 2C) and flow cytometry (Figure 2D). We noticed that over time in culture the proportion of Xmas ESCs that were single positive for the fluorescent reporters progressively increased, likely reflecting increasing abundance of XO cells. Therefore, we tested whether the fluorescent reporters accurately reflected karyotype. We used fluorescence activated cell sorting (FACS) followed by PCR of genomic DNA and found the GFP reporter allele detectable only in the GFP^+^ population, the mCherry reporter only in the mCherry^+^ population and both reporter alleles in the double positive population (Figure S2A). Thus, the reporter alleles detect XX and XO cell populations, while also accurately reflecting the XCI status in XX cells.

**Figure 2.**
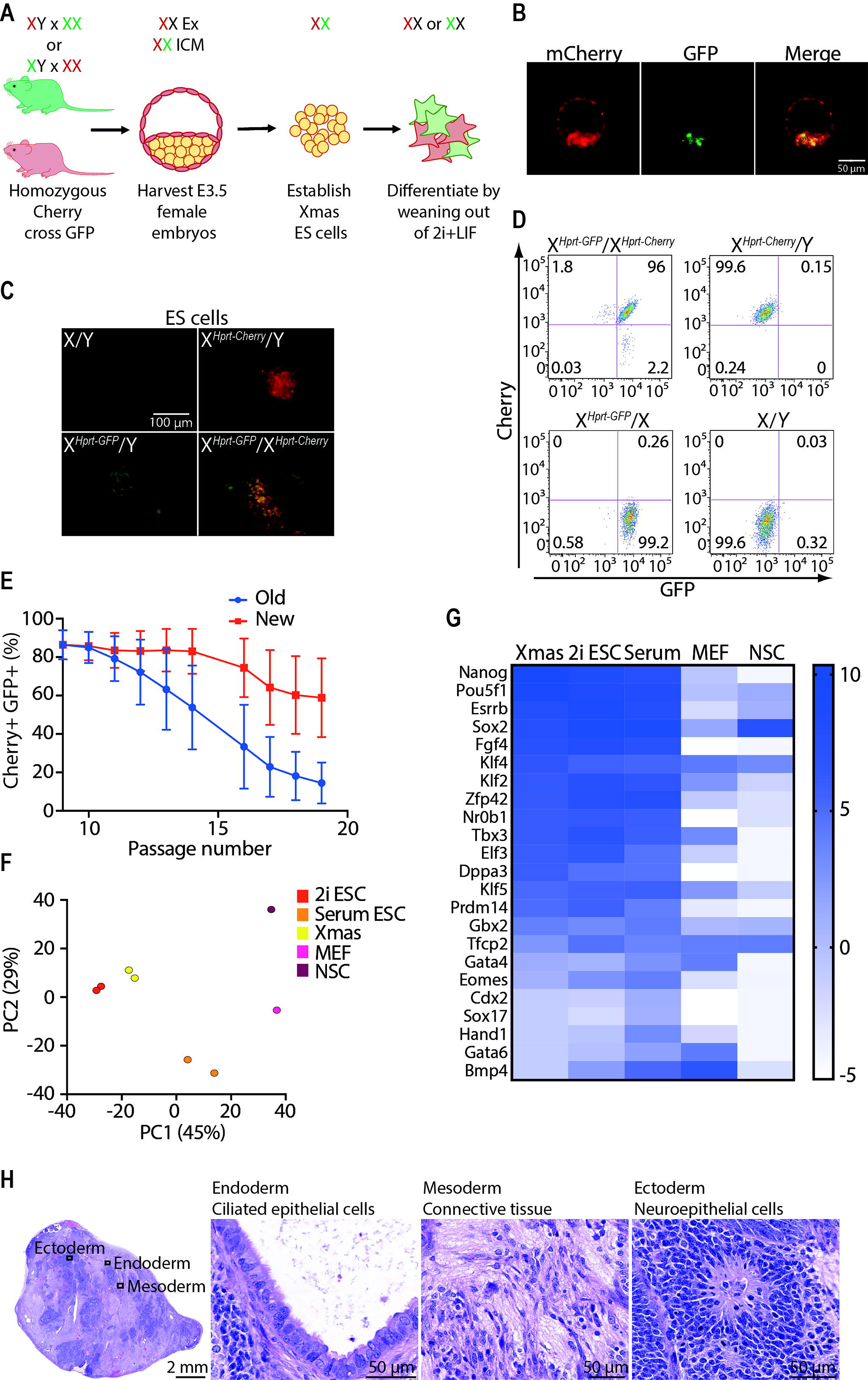
Xmas ESCs enable development of improved culture conditions. **(A)** Schematic of the breeding strategy to produce Xmas ESCs, their XCI status *in vivo* and during culture and differentiation *in vitro*. Extraembryonic (Ex), Inner Cell Mass (ICM) **(B)** Live fluorescent image of X^*Hprt-*GFP^/X^*Hprt*-mCherry^ Xmas female blastocysts. **(C)** Live fluorescent image of cultured ESCs from the indicated genotypes carrying different combinations of the fluorescent reporter alleles. **(D)** Flow cytometry of cultured ESCs from the indicated genotypes carrying different combinations of the fluorescent reporter alleles. **(E)** Flow cytometry data from primary female ESCs maintained in 2i media over 18 passages in either traditional (Old) or our improved (New) culture conditions, where presence of the reporter alleles indicates the X karyotype of the cell. **(F)** Principle component analysis of RNA-seq data from Xmas ESCs compared to published transcriptomes of ESCs grown in serum or 2i, MEFs or neural stem cells (NSCs). (**G)** Heat map showing average expression (log_2_ rpm) of pluripotency genes in Xmas ESCs (n = 2) and published transcriptomes of ESCs grown in serum or 2i, MEFs or NSCs. **(H)** Representative images of teratomas produced following injection of Xmas ESCs into nude mice (n = 4), with differentiated cell types from endodermal, mesodermal and ectodermal lineages shown.

The ease with which we could monitor the presence of the X chromosomes in Xmas ESCs allowed us to assess and improve current methods for the maintenance of female ESCs. Testing of different parameters lead to the identification of conditions that substantially improve maintenance of the XX karyotype (Figure 2E), with the major improvements being made through the use of 2i media (Ying et al., 2008), daily passaging and the lack of an attachment substrate (Figure S2B). We provide the full optimised protocol as part of the methods. To further characterise Xmas ESCs we performed RNA-seq and compared their transcriptomes to those of previously published ESCs (Marks et al., 2012; Maza et al., 2015). Principle component analysis showed that they most closely resembled ESCs grown in 2i media (Figure 2F), with similar expression of pluripotency genes (Figure 2G). These data suggest that derivation and culture of Xmas ESCs under our improved culture conditions do not alter their naïve pluripotent state (Ying et al., 2008). Finally, we tested whether Xmas ESCs made via our new protocol retain pluripotency by performing a teratoma formation assay. We found unconstrained formation of teratomas that displayed differentiation into all three germ layers (Figure 2H). These data confirm that Xmas ESCs produced via our protocol are pluripotent cells in the naïve state, as expected, and are therefore a useful tool with which to study female pluripotency.

### Refined culture system improves male ESC fitness

We next asked whether our method was also beneficial for the maintenance of male ESCs. We derived two lines of wild-type male ESCs on the C57BL/6 background and cultured them either using our protocol (daily passaging in 2i, no attachment substrate) or the current state-of-the-art ESC culture protocol (passaging every 2 days in 2i media, with gelatin attachment substrate) (Mulas et al., 2019), taking samples at passages (p) 0, 10 and 20 and performing both RNA-seq and DNA-seq. Multi-dimensional scaling analysis of the RNA-seq data showed that our improved protocol maintained the male ESCs in a transcriptionally similar state to freshly derived p0 cells for both ESC lines, whereas cells maintained under traditional conditions diverged significantly at both p10 and p20 (Figure 3A), suggesting our method was keeping the cells transcriptionally closer to the starting population. We identified 5526 differentially expressed genes (False discovery rate (FDR) < 0.05) between the two culture methods when p10 and p20 were combined, with gene set testing identifying ribosome and mitochondrial associated genes as being significantly upregulated in cells maintained under our improved conditions (Figure 3B,C, Table S1), consistent with these cells being in a rapid state of self-renewal.

**Figure 3.**
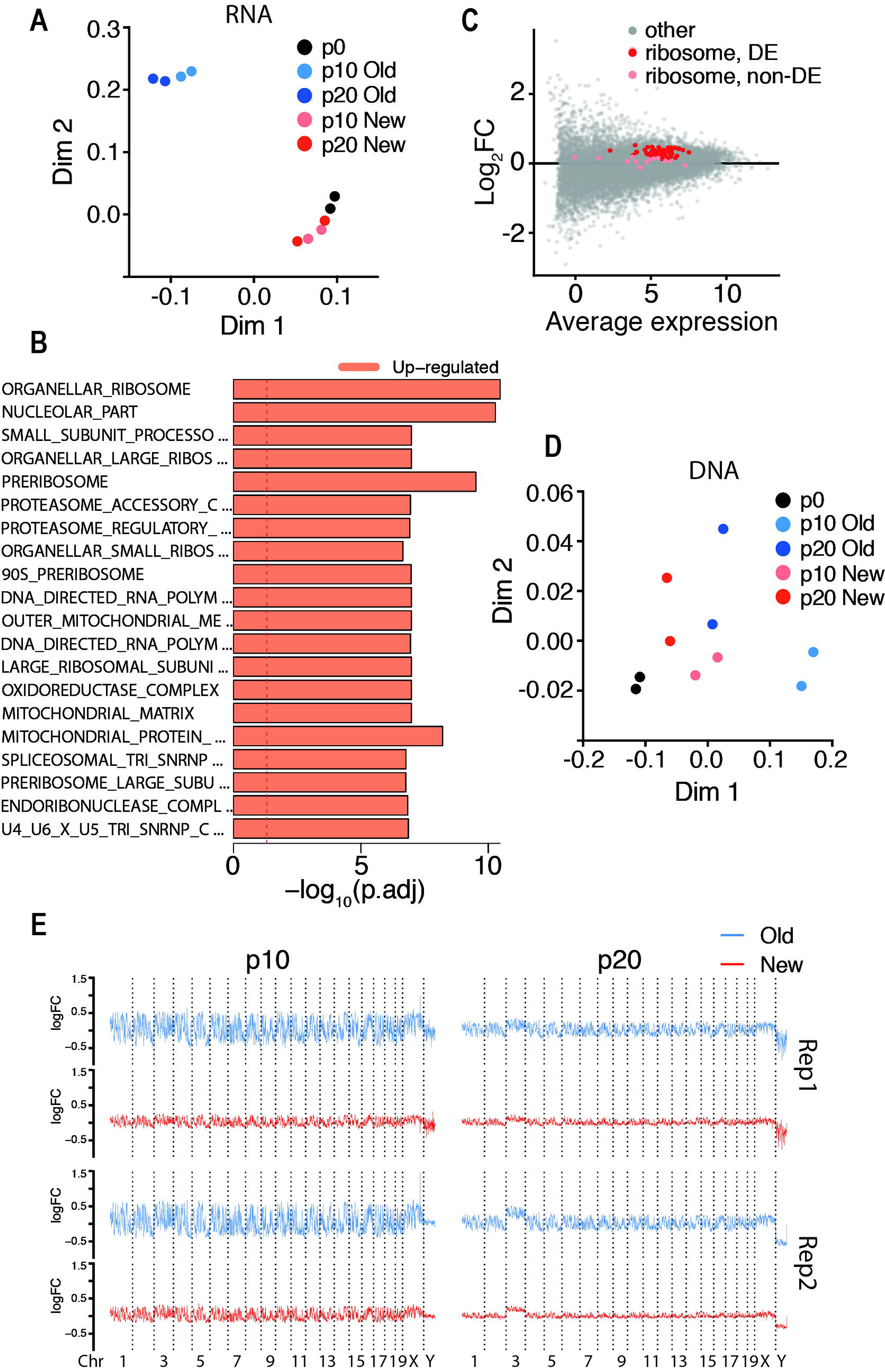
New culture conditions maintain transcriptome and karyotype of male ESCs. **(A)** Multi-dimensional scaling (MDS) plot of RNA-seq data from p0 male ESCs compared to cells cultured in either old or new culture conditions at p10 and p20 (n = 2). **(B)** Results of gene set testing from combined p10 and p20 samples using either the old or new culture conditions. Dashed line indicates p-value = 0.05. **(C)** MA plot showing the average fold change (log_2_FC) at p10 and p20 combined of genes between male ESCs grown in our new vs old culture conditions. Significantly differentially expressed ribosomal genes are indicated in red and non-significant ribosomal genes in pink. **(D)** MDS plot of DNA-seq data in 1Mb bins from p0 male ESCs compared to cells cultured in either old or new culture conditions at p10 and p20 (n = 2). **(E)** Read coverage plots of 1 Mb bins across all chromosomes of male ESCs cultured in either old or new conditions with reads normalised to equivalent positions in respective p0 samples from two replicate cell lines.

To assess the effect of the two culture methods on the karyotype and copy number variation of male ESCs we analysed our DNA-seq data and found that one of the two cell lines had lost the Y chromosome prior to p0, while the second cell line was depleted by p20. This occurred in both culture methods and is consistent with the Y chromosome being dispensable for male cells *in vitro* (Figure S3). Principle component analysis showed that DNA obtained from cells cultured under our improved conditions for 10 and 20 passages was most similar to that from p0, where again cells cultured with the traditional method diverged (Figure 3D); consistent with our method maintaining a karyotype most similar to the starting population. We next sought to identify what the defining differences in the DNA were by analysing the genome in 1Mb bins. At p10, we found that when cultured by our new method none of the genome was differentially represented (FDR < 0.05, log_2_FC > 1.1), while cells cultured by the traditional method had ~57% of the genome differentially represented (Figure 3E, Table S2). At p20, cells cultured by the new and old methods had 0% and ~5.5% differentially represented regions respectively, suggesting that prolonged culture selected against karyotypic abnormalities. Taken together, these data indicate that our method of culture maintains male ESCs transcriptionally and karyotypically in a state that most closely resembles freshly derived ESCs and that our method can be applied to better maintain both male and female ESCs *in vitro*.

### Xmas reporter alleles detect iPSC reprogramming

Xmas ESCs are a highly tractable system for the study of female ESCs, therefore we next sought to assess their utility for studies of female pluripotency more broadly. Given that reactivation of the Xi is seen as a key indicator of cellular reprogramming (Pasque et al., 2014), we reasoned that our reporter alleles may also be useful to study induced pluripotent stem cell (iPSC) generation. We crossed X^*Hprt*-GFP^ and X^*Hprt*-mCherry^ mice and derived post-XCI Xmas MEFs from the resulting E13.5 embryos. As before (Figure 1F), we found GFP^+^ and mCherry^+^ cells in roughly equal numbers, with no MEFs found to be double positive, indicating complete XCI as expected. We purified the GFP^+^ or mCherry^+^ populations by FACS and transduced them with lentiviral vectors containing a doxycycline inducible reprogramming cassette (STEMCCA) (Sommer et al., 2009) which encodes the transcription factors *Oct4, Klf4, Sox2* and *c-Myc* (OKSM) and a reverse tetracycline transactivator (rtTA). Reprogramming was induced at day 0 by the addition of doxycycline and cells were monitored by flow cytometry for 16 days (Figure 4A). As expected, cells were completely single positive for the fluorescent markers at day 0 of reprogramming, remaining so until day 12 when the first double positive cells appeared, with ~80% of cells expressing both markers by the end of the assay (Figure 4B), indicating reactivation of the Xi and further confirming that reactivation of the Xi is a late event during reprogramming. We are unable to say whether the ~20% of cells that did not become double positive were due to a failure to reactivate the Xi or due to loss of an X chromosome. To determine if iPSCs are of unstable karyotype, similar to ESCs, we maintained our iPSCs through a number of passages using regular iPSC culture techniques, assessing karyotype by flow cytometry. Stunningly, iPSCs became XO much faster than ESCs, with only ~2% of cells remaining double positive by passage 6 (Figure 4C). These data highlight a critical need for improved methods to maintain karyotype in female iPSCs *in vitro*.

**Figure 4.**
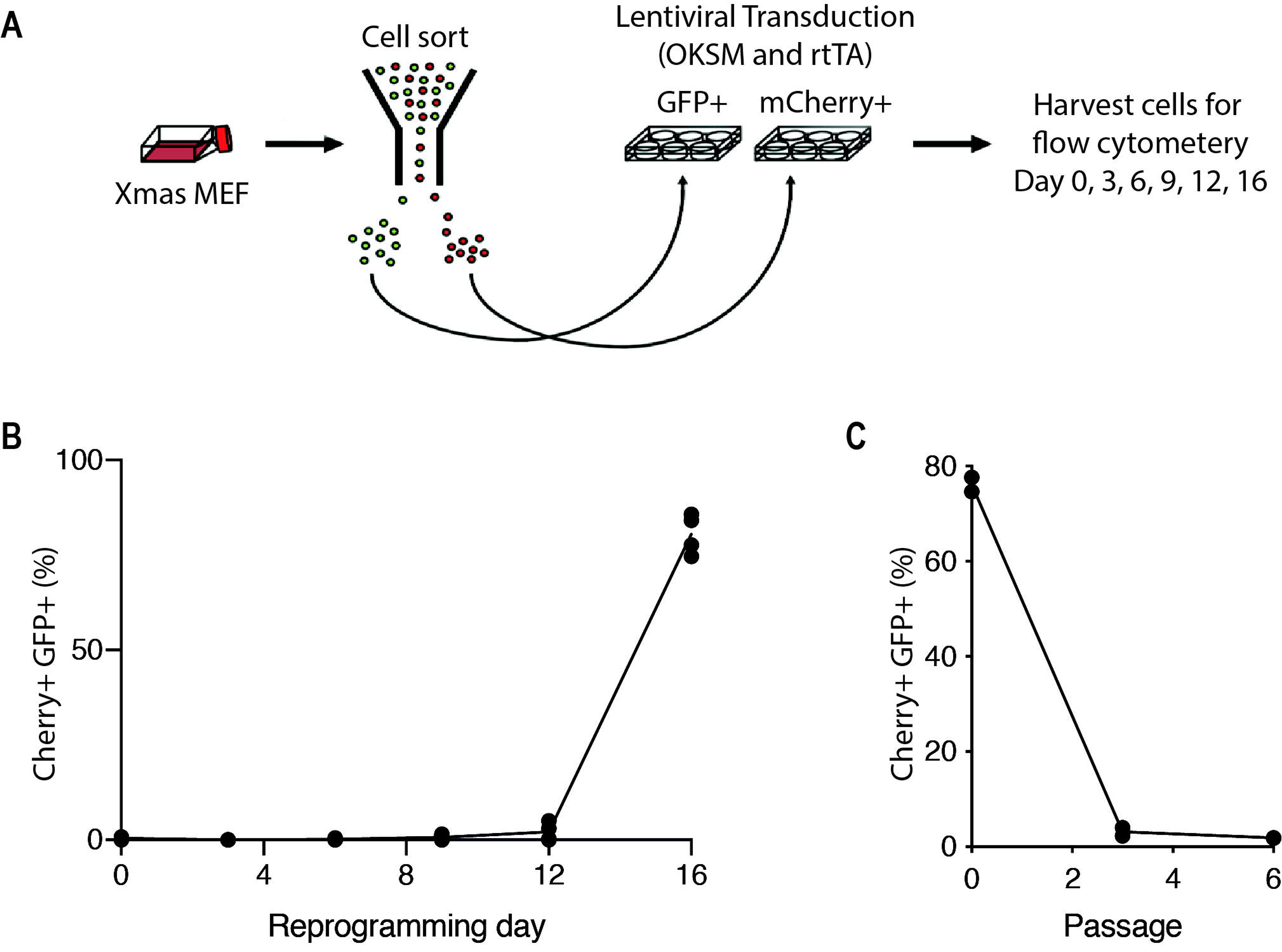
Xmas reporter alleles detect Xi reactivation during iPSC reprogramming. **(A)** Schematic showing the strategy for reprogramming and analysis of Xmas MEFs. **(B)** Flow cytometry data from primary female Xmas MEFs during the reprogramming process (n = 4). **(C)** Flow cytometry data from reprogrammed Xmas iPSCs in standard iPSC maintenance media for 6 passages (n = 2).

### Xmas ESCs have the capacity to detect impaired XCI

Xmas ESCs, together with our improved culture conditions, allowed us to consider functional studies of XCI. We first asked whether our reporter alleles could detect random XCI upon differentiation (Figure 2A). Xmas ESCs were induced to differentiate on day 0 by weaning them out of 2i media and into differentiation media over 3 days, with the cells monitored daily by flow cytometry. At day 0, the majority of all cell lines tested were predominantly double positive for the fluorescent markers, indicating an XaXa XCI state. From days 5 through to 7 of differentiation there was a rapid loss of double positivity with all cell lines tested adopting the XaXi state expected following random XCI (Figure 5A). To determine whether differentiation was proceeding normally in these cells, we performed RNA-seq along the same time course and compared this to published datasets (Marks et al., 2012; Maza et al., 2015). As expected, Xmas ESCs begin differentiation most closely transcriptionally aligned to the naïve state before transitioning through a state similar to primed pluripotency and finally most closely resembling MEFs by day 8 of differentiation (Figure 5B, Table S3). Therefore, Xmas ESCs differentiated normally and showed at the single cell level the timing of XCI during differentiation.

**Figure 5.**
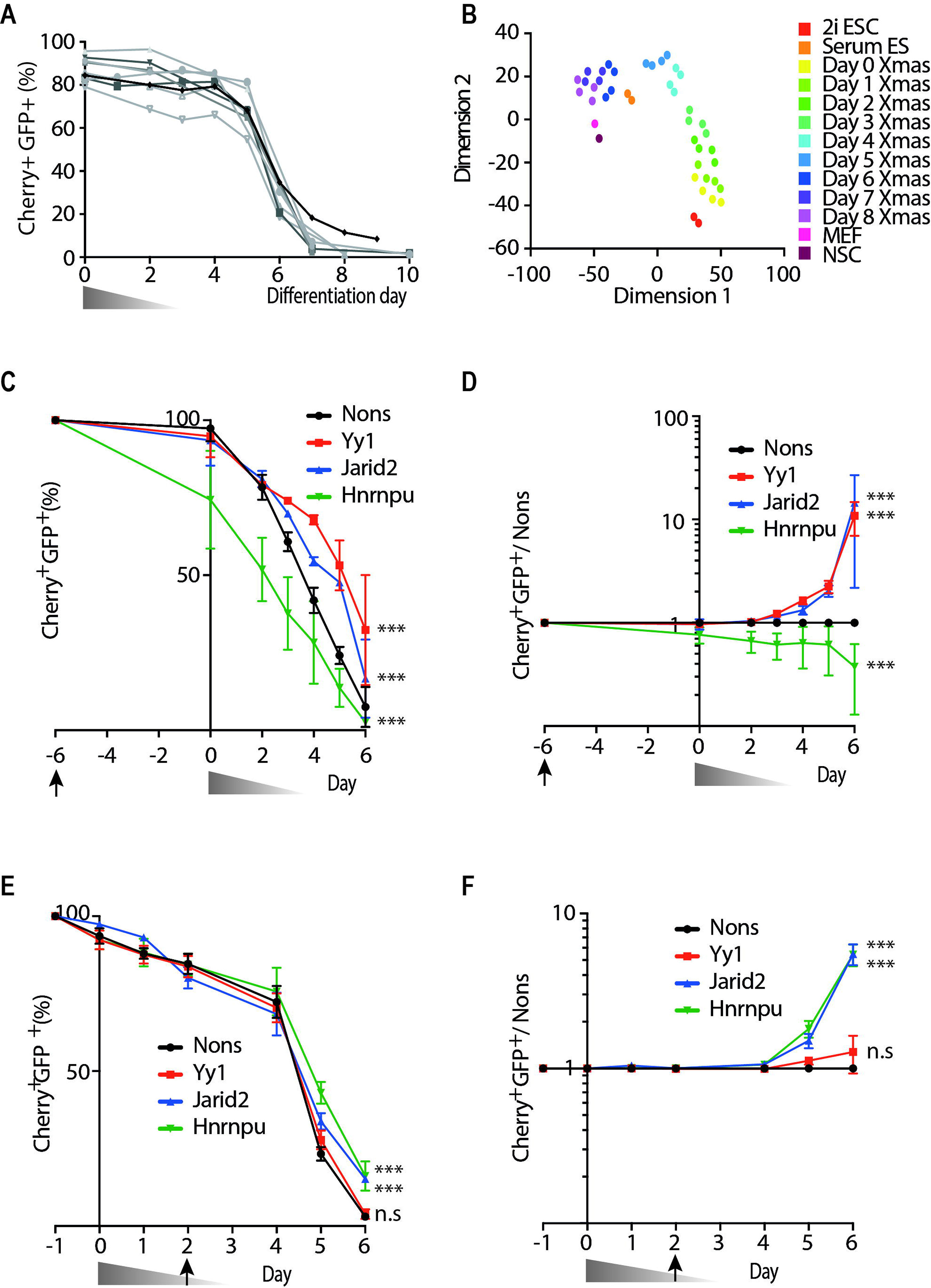
Xmas ESCs detect impaired XCI during differentiation. **(A)** Flow cytometry data showing the kinetics of the fluorescent reporter alleles during differentiation and XCI of Xmas ESCs for multiple cell lines (n = 9). The triangle represents weaning from 2i to differentiation media in 25% increments over 3 days. **(B)** tSNE plot comparing the transcriptomes of Xmas ESCs (n = 4) from day 0 to day 8 of differentiation against published transcriptomes of ESCs grown in serum or 2i, MEFs or NSCs. **(C-F)** Flow cytometry data showing the kinetics of the fluorescent reporter allele expression changes during differentiation and XCI of Xmas ESCs. Cells were challenged with shRNAs against the indicated known regulators of XCI or control (Nons) either prior to differentiation depicted as either raw data **(C)** or normalised to Nons **(D)**, or during differentiation as either raw data **(E)** or normalised to Nons **(F)**. Triangles represent weaning from 2i media into differentiation media and arrows indicate the day of shRNA viral transduction. n = 3–5 for each of two independent shRNAs per gene, error bars indicate s.e.m., one-way ANOVA, *** indicates *p* < 0.001.

We next tested if XCI could be perturbed in Xmas ESCs by knockdown of genes known to regulate XCI, including Yy1 which is required for the transcriptional initiation of *Xist* (Makhlouf et al., 2014), Hnrnpu which helps tether *Xist* to the future Xi (Hasegawa et al., 2010) and Jarid2 which directs polycomb-mediated repression to *Xist*-localised regions (Cooper et al., 2016; da Rocha et al., 2014). We transduced Xmas ESCs with virus containing validated shRNAs against *Yy1*, *Hnrnpu* and *Jarid2* (Figure S4) 6 days prior to differentiation and monitored XCI by flow cytometry. We found that both knockdown of *Yy1* and *Jarid2* were able to inhibit XCI relative to a non-silencing negative control (Nons) (Figure 5C). To overcome the variable percentage of XO cells in the starting populations, each experiment was normalised to the matched Nons control (Figure 5D). Unexpectedly, knockdown of *Hnrnpu* caused a rapid loss of double positivity prior to weaning into differentiation media, likely due to the requirement of Hnrnpu for pluripotency (Vizlin-Hodzic et al., 2011) and suggesting Xmas ESCs may be useful for the identification of novel pluripotency factors. To circumvent the role of Hnrnpu in pluripotency and instead test its function in XCI, we transduced cells with shRNA at day 2 of differentiation, so that knockdown takes effect following exit from pluripotency. Using this strategy, *Hnrnpu* knockdown no longer caused accelerated loss of double positivity, but rather the expected inhibition of XCI relative to the control (Figure 5E,F). However, depletion of *Yy1* at this timepoint no longer delayed XCI, despite *Jarid2* knockdown continuing to have an effect. This result is consistent with the role of Yy1 early in the process of XCI (Donohoe et al., 2007; Makhlouf et al., 2014) prior to when knockdown is achieved if cells are transduced at day 2 of differentiation. These data suggest that by varying the time of shRNA transduction, Xmas ESCs may be used to dissect the stage of random XCI for which each factor is required and therefore they may provide a deeper understanding of the kinetics of the process of random XCI.

### Smarcc1 and Smarca4 are required for XCI

As Xmas ESCs permitted us to grow female ESCs with two X chromosomes for longer than before, and because Xmas ESCs could readily be used to detect the effects of known regulators of XCI, we next performed a screen aimed at detecting new genes required for the establishment of XCI. Our previous mouse genetic screen was successful in identifying epigenetic regulators of transgene variegation (Ashe et al., 2008; Blewitt and Whitelaw, 2013; Blewitt et al., 2005; Chong et al., 2007; Daxinger et al., 2013; Daxinger et al., 2012; Harten et al., 2014; Whitelaw et al., 2010; Youngson et al., 2013). We and others have shown that several of the novel and known epigenetic regulators identified in this screen are also required for XCI, including Smchd1, Setdb1 and Dnmt1 (Blewitt et al., 2008; Keniry et al., 2016; Minajigi et al., 2015; Minkovsky et al., 2014; Sado et al., 2000). Given this, we decided to screen the suite of genes that emerged from the mouse genetic screen for roles in XCI using our Xmas ESC system, to try and identify new regulators of XCI. Xmas ESCs were transduced on day 2 of differentiation with validated hairpins against candidate genes (Figure S4) and assessed by flow cytometry at day 6; a timepoint at which we consistently observe an effect of gene knockdown on XCI. Prominently, two independent shRNAs that knockdown expression of *Smarcc1* and *Smarca4* caused a substantial impairment of XCI (Figure 6A). By contrast, knockdown of known regulators of the maintenance phase of XCI (*Dnmt1*, *Smchd1*) did not produce a readout in this screen, suggesting day 6 of differentiation is too early in the timecourse of XCI to reveal factors that have a role exclusively in the maintenance phase of silencing. We validated this result in a Xmas ESC differentiation time course, where we found knockdown of both *Smarcc1* and *Smarca4* led to a measurable increase in double positive cells by day 5 of differentiation (Figure 6B and S5A). This is the first screen for the establishment of XCI performed in near-native female ESC, made possible via the Xmas ESC system and our improved culture conditions.

**Figure 6.**
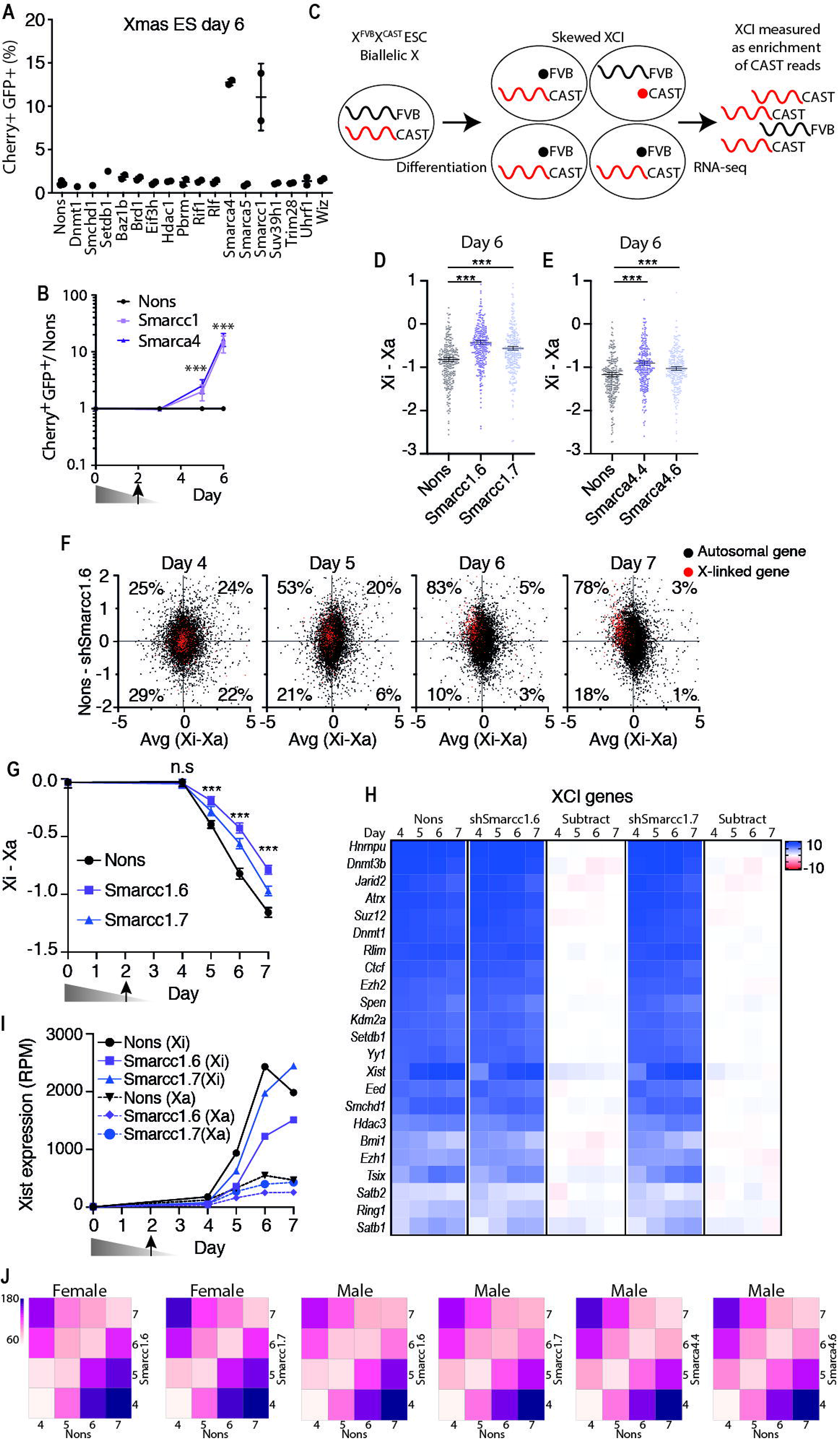
Screen in Xmas ESCs identifies Smarcc1 and Smarca4 as regulators of XCI. **(A)** Flow cytometry data at day 6 of Xmas ESC differentiation following viral transduction of shRNAs at day 2 against candidate genes (n = 2 independent hairpins per gene, error bars indicate S.D.). **(B)** Flow cytometry data normalised to Nons along a time course of Xmas ESC differentiation following shRNA mediated knockdown of *Smarcc1*, *Smarca4* or Nons (n = 4 for each of two independent shRNAs per gene, error bars indicate s.e.m., Student’s paired *t*-test, *** indicates *p* < 0.001). **(C)** Schematic of skewed XCI during differentiation of X^FVB^X^CAST^ ESCs. **(D,E)** Allele-specific RNA-seq data of X^FVB^X^CAST^ ESCs at day 6 of differentiation following knockdown with indicated hairpins against *Smarcc1* **(D)** and *Smarca4* **(E)**. Each point represents the Xi-Xa log_2_ expression value of an individual informative X-linked gene (error bars indicate s.e.m., Student’s paired *t*-test, *** indicates *p* < 0.001). **(F)** These graphs show RNA-seq data and are designed to compare gene expression from the X chromosome to autosomes. Each point on the graph represents an informative gene, with X-linked genes in red and autosomal genes in black. The x-axis shows the ratio of expression from FVB compared to CAST (Xi − Xa log_2_), therefore XCI is observed as a left shift of the red dots along the x-axis. The y-axis shows the ratio of expression from Nons compared to knockdown with Smarcc1.6 (Nons − Smarcc1.6 log_2_FC), therefore a failure of XCI in the knockdown is observed as an upward shift of the red dots along the y-axis. Black dots give an indication of global trends in autosomal gene expression. Dotted lines indicate medians and percentages show the X-linked genes falling into each quadrant. (**G)** RNA-seq time course data showing the ratio of Xi gene expression compared to the Xa (Xi – Xa log_2_). Error bars show the s.e.m. of all informative genes, Student’s paired *t*-test, *** indicates *p* < 0.001. **(H)** Heat map of gene expression (rpm log_2_) of known regulators of XCI, with the difference between knockdown and control (subtract, Nons − knockdown) indicated. **(I)** Expression of *Xist* (rpm log_2_). Triangle represents weaning from 2i media into differentiation media and arrows indicate the day of shRNA viral transduction. **(J)** Heat maps showing the average Euclidean distance in gene expression (log_2_cpm) between knockdown and Nons control along a differentiation time course of either male or female ESCs.

To determine the extent to which *Smarcc1* and *Smarca4* knockdown impaired XCI, we performed allele-specific RNA-seq. We derived wild-type F1 ESCs by crossing FVB/NJ (FVB) dams with CAST/EiJ (CAST) sires. These ESCs were cultured using our improved conditions to maintain karyotype, before being differentiated and transduced with shRNA at day 2 of differentiation. X^FVB^X^CAST^ ESCs allow for allele-specific analyses of XCI as the parental alleles can be discriminated by single nucleotide polymorphisms (SNPs) and XCI is naturally skewed, with the FVB allele approximately 3 times more likely to become the Xi and the CAST to become the Xa upon establishment of XCI (Figure 6C, S5B,C). Naturally skewed cells avoid the need to genetically skew random XCI by deletion of *Xist,* thereby allowing the normal process of XCI to occur. For simplicity, we refer to the X^FVB^ as the Xi and the X^CAST^ as the Xa. Both *Smarcc1* and *Smarca4* knockdown resulted in an increase in gene expression from the Xi at day 6 of differentiation at the majority of informative X-linked genes (Figure 6D,E, S5B–D, Table S4,5), suggesting Smarcc1 and Smarca4 are both required for chromosome-wide silencing. To gain a more detailed insight into the kinetics of silencing we focussed on *Smarcc1* knockdown and performed RNA-seq along a differentiation time course. We found a persistent failure of XCI in *Smarcc1* knockdown cells compared to control that was detectable from day 5, while no increase in gene expression from the future Xi (the FVB allele) was detectable prior to the onset of XCI at day 4 (Figure 6F,G, S5E).

We next reviewed data from the RNA-seq time course for possible causes of impaired XCI upon *Smarcc1* knockdown. There were no significantly differentially expressed genes in common between *Smarcc1* and *Smarca4* knockdown groups, suggesting the mechanism by which Smarcc1 and Smarca4 regulate XCI is not via a secondary gene or delayed differentiation. Consistent with this interpretation, we found no substantial deregulation of genes known to be involved in XCI upon either *Smarcc1* (Figure 6H) or *Smarca4* depletion (Figure S5G). There was also no difference in the timing of *Xist* induction upon *Smarcc1* knockdown compared to controls suggesting that XCI is initiated correctly, although the expression level of *Xist* was slightly reduced (Figure 6H,I). To assess whether *Smarcc1* and *Smarca4* knockdowns were causing delayed differentiation, we analysed the transcript kinetics of our female differentiation RNA-seq time course as well as a time course in male cells, finding no consistent delay in differentiation upon depletion of these nucleosome remodelling factors (Figure 6J, Table S6). Finally, to experimentally separate the role of Smarcc1 and Smarca4 on XCI from their role in pluripotency, we differentiated Xmas ESCs and transduced them with shRNA at day 3, such that knockdown is achieved at approximately day 4 and found that both *Smarcc1* and *Smarca4* knockdown still caused a detectable failure of XCI (Figure S5F). Taken together, these data suggest that Smarcc1 and Smarca4 have a direct effect on the establishment of X chromosome silencing, as opposed to delaying differentiation or as potential regulators of *Xist* or other known XCI regulators.

### Smarcc1 and Smarca4 are required at the establishment phase of XCI

We next undertook a series of experiments to test our interpretation of the RNA-seq data, that suggested Smarcc1 and Smarca4 were required for the establishment phase of XCI. Firstly, to exclude a role for Smarcc1 and Smarca4 in the initiation of XCI we performed RNA fluorescence *in situ* hybridisation (FISH) for *Xist* RNA in differentiating ESCs at day 4 and 5. We found no difference in the number of cells with an *Xist* focus at either day between knockdown of *Smarcc1* or *Smarca4* and controls (Figure 7A,B, S6A,B). Importantly, there was a substantial increase in the number of cells with *Xist* foci between days 4 and 5, indicating that these cells were captured at a time when they were initiating XCI. These data exclude a role of Smarcc1 and Smarca4 in the initiation of XCI.

**Figure 7.**
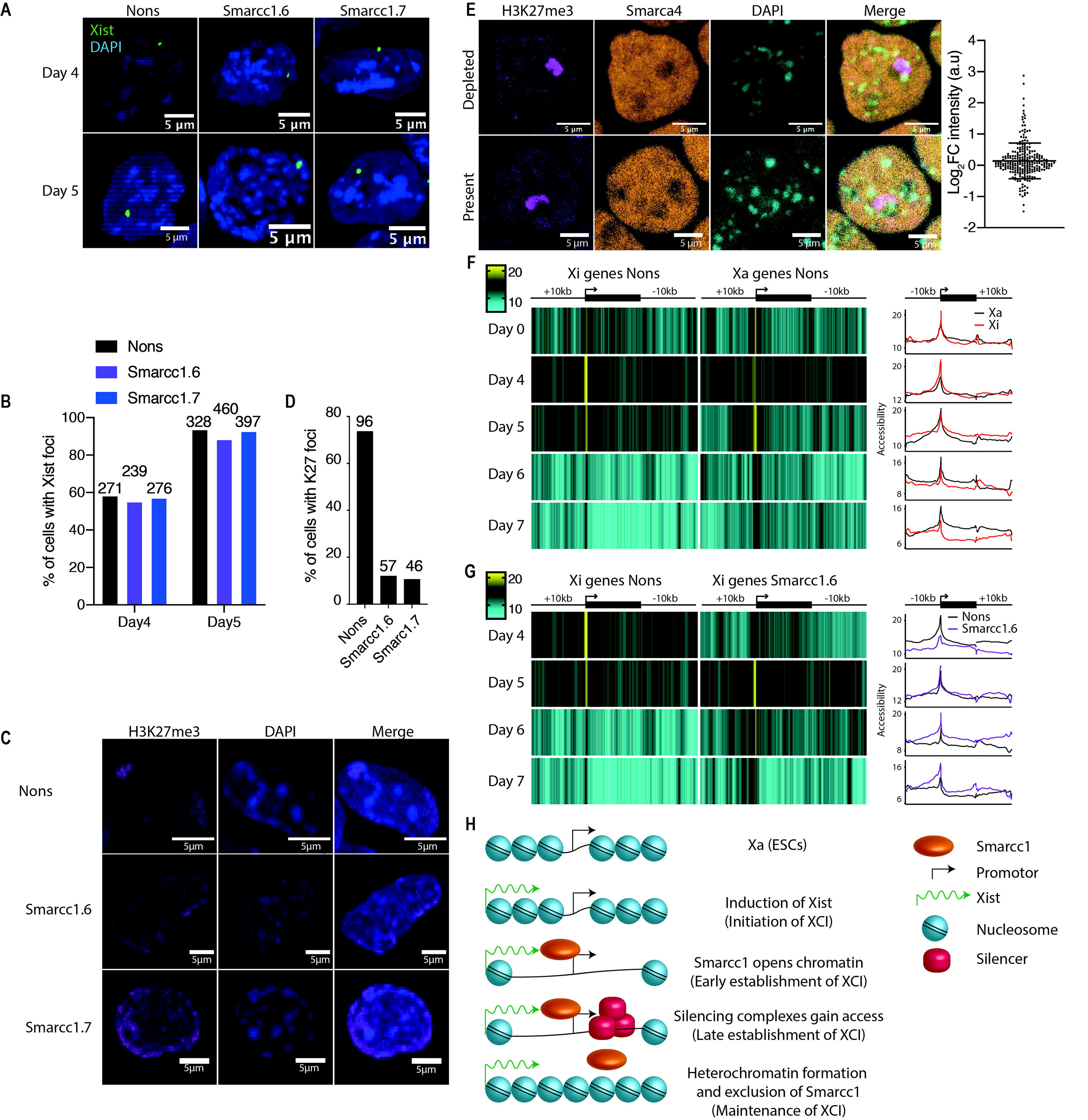
Smarcc1 opens Xi promotors in order for establishment of XCI to proceed. **(A)** RNA FISH for *Xist* in female ESCs at day 4 and 5 of differentiation following knockdown with the indicated hairpins. **(B)** Quantification of data from (A). **(C)** Immunostaining of H3K27me3 in female ESCs at day 6 of differentiation following knockdown with the indicated hairpins. **(D)** Quantification of data from (C). **(E)** Immunostaining of H3K27me3 and Smarca4 in female ESCs at day 6 of differentiation. Representative images of Smarca4 being both present and depleted at sites of H3K27me3 are shown. Plot indicates the average intensity in arbitrary units (a.u) underneath H3K27me3 foci compared to the rest of the cell (log_2_FC), for all cells measured. **(F,G)** Nucleosome occupancy (% GpC methylation) along a time course of female ESC differentiation determined by NOMe-seq averaged across all genes and flanking regions on the Xi and Xa **(F)** or the Xi upon *Smarcc1* knockdown **(G)**. The data shown as a heat map or a smoothed histogram. **(H)** Model for how Smarcc1 regulates establishment of XCI.

To test the role of Smarcc1 and Smarca4 in the establishment of XCI we performed immunofluorescence for H3K27me3, a marker of an established Xi, in differentiating ESCs at day 6 of differentiation. We found that both *Smarcc1* and *Smarca4* knockdown caused a substantial reduction in cells with a H3K27me3 focus compared to the control (Figure 7C,D, S6C,D). These data are suggestive of a role for both Smarcc1 and Smarca4 in the establishment of XCI, however they do not rule out a role in the maintenance of XCI. To test Smarcc1 and Smarca4 in the maintenance phase of XCI we performed knockdown experiments in post-XCI Xmas MEFs and analysed them by flow cytometry following treatment with the DNA methylation inhibitor 5-azacytidine. We found knockdown of either gene was unable to reactivate the silent reporter allele (Figure S7A). As the reversal of XCI in the maintenance phase is difficult, we next turned to a more sensitive reporter system (Csankovszki et al., 2001; Graves, 1982; Hadjantonakis et al., 2001; Hadjantonakis et al., 1998; Mohandas et al., 1981), where MEFs carry a silent multi-copy GFP transgene on their Xi by virtue of a *Xist* knockout allele *in trans* to the reporter (Xi^GFP^Xa^Δ*Xist*^ MEFs) (Keniry et al., 2016; Royce-Tolland et al., 2010). Again, we found no reactivation of the GFP reporter upon either *Smarcc1* or *Smarca4* knockdown, despite our *Dnmt1*, *Smchd1* and *Setdb1* knockdown positive controls producing readily detectable GFP expression (Figure S7B). Moreover, by using H3K27me3 enrichment as a marker of the inactive X at day 6 of ESC differentiation, we were able to test whether Smarca4 was present on the Xi by immunofluorescence. We found that Smarca4 was present on the Xi in some cells but depleted in others, suggesting the Xi has a heterogeneous requirement for Smarca4 at this early stage of differentiation. As Smarca4 is known to be absent from the Xi in terminally differentiated cells (Jegu et al., 2019; Minajigi et al., 2015), these data are consistent with Smarca4 being present on the Xi during establishment of XCI, but excluded upon completion of XCI (Figure 7E, S6D). Taken together these data provide strong evidence that both Smarcc1 and Smarca4 are required to establish, rather than to initiate or maintain, XCI. This is the first report of this role for Smarcc1 and Smarca4 in the establishment of XCI.

### Smarcc1 depletes nucleosome occupancy at Xi promotors prior to establishment of silencing

The fact that both Smarcc1 and Smarca4 contribute to the establishment phase of XCI, suggested that they may function directly on the Xi through their enzymatic capacity as nucleosome remodellers. To test this, we profiled nucleosome occupancy in differentiating X^FVB^X^CAST^ ESCs by allele-specific Nucleosome Occupancy and Methylome Sequencing (NOMe-seq) (Kelly et al., 2012; Lay et al., 2018; Taberlay et al., 2014). The use of this technique to study the establishment of XCI has not been reported previously, so we initially concentrated on the normal time course of XCI in ESC differentiation (Nons control). The reduced coverage provided by allele-specific data precluded gene-specific analyses, however we were able to profile XCI by averaging across all X-linked genes, finding different nucleosome remodelling kinetics between the Xi and Xa. On the Xa, promotors are slightly open in ESCs, remaining similarly open at day 4 of differentiation, then opening further at day 5, before restricting again at day 6 (Figure 7F). As expected, the Xa kinetics were mirrored on the autosomes (Figure S6F), and show promotors becoming more open as cells transition from pluripotency to a more lineage restricted state. Strikingly, the Xi follows a different pattern of nucleosome remodelling, and we know that the vast majority of the Xi genes will undergo silencing in concert on the following days (Figure 6F,G, S5B,E). For the Xi, promotors are initially slightly open, similarly to those of the Xa, but become more nucleosome-depleted at day 4 of differentiation, a day earlier than the depletion observed on the Xa (Figure 7F). These data suggest that promotors of the Xi become accessible prior to establishment of XCI. Following day 4, the Xi becomes progressively more heterochromatic with both promotors and gene bodies becoming increasingly nucleosome dense.

To address the functional role of nucleosome depletion prior to gene silencing, we utilised a *Smarcc1* depleted NOMe-seq time course, as we had matched RNA-seq data in X^FVB^X^CAST^ ESC differentiation. Remarkably, *Smarcc1* depleted cells were unable to open promotors at day 4 on the Xi and instead followed similar kinetics to the Xa, consistent with the failure of gene silencing we observed in our RNA-seq data. These data suggest that an inability to open promotors at day 4 results in a failure to establish XCI (Figure 7G), meaning that the nucleosome depletion of promoters on the Xi at day 4 is required for subsequent gene silencing. We note that our genomic data obtained from X^FVB^X^CAST^ ESCs hides the full magnitude of the effects, as they display partially skewed rather than completely skewed XCI. No significant effect of *Smarcc1* depletion was observed at a chromosome-wide level on the Xa (Figure S6E) or at a genome-wide level on autosomes (Figure S6F), however there are likely to be gene-specific abnormalities that we did not have the power to detect with NOMe-seq and undirected differentiation. An advantage of the NOMe-seq method is that it also provides information on methyl-cytosine. As expected, ESCs were globally hypomethylated, remaining hypomethylated at gene promotors during differentiation, but becoming increasingly more methylated at intergenic regions and gene bodies. The methylation of CpG island promoters on the Xi is a feature of the maintenance phase of XCI; as expected given the timing of our samples, we did not observe such methylation occurring in our time course, and there was no difference observed between the Xi and Xa (Figure S6G,H). Therefore, we have identified a new role for chromatin remodeller Smarcc1 in the earliest stages of the establishment of XCI and exit from pluripotency and provide the first example of chromatin opening being a necessary initial step towards gene silencing.

## Discussion

Female pluripotency is different to male pluripotency. Indeed, female pre-implantation embryos are found to develop more slowly than their male counterparts, with this difference being attributed to the dosage imbalance of the sex chromosomes that occurs during this developmental period (Burgoyne et al., 1995; Gardner et al., 2010; Mittwoch, 1993; Schulz, 2017). Embryonic stem cells provide a tractable *in vitro* model in which to study the pluripotency of the pre-implantation embryo. These cells exhibit many of the traits associated with the cells of the inner cell mass from which they are derived, including activity from both parental X chromosomes in females. Double dosage of X-linked genes has been found to increase pluripotency factor expression, while also inhibiting targets of the differentiation promoting Mek/Erk signalling pathway (Choi et al., 2017a; Schulz et al., 2014), having the combined effect of driving female ESCs further towards a ground state of pluripotency compared to males. Moreover, female ESCs are delayed in their exit from pluripotency upon differentiation *in vitro*, only acquiring a differentiated transcriptome similar to male cells following complete XCI (Chen et al., 2016; Schulz et al., 2014). Ground state pluripotency is associated with low levels of DNA methylation and consistent with this, female ESCs display greatly reduced levels compared to those seen in males at all genomic features (Choi et al., 2017a; Choi et al., 2017b; Habibi et al., 2013; Ooi et al., 2010; Schulz et al., 2014; Yagi et al., 2017; Zvetkova et al., 2005). This effect is known to be dependent on dosage of the X-linked gene *Dusp9* (Choi et al., 2017a). Strikingly, XO female ESCs resemble XY males both functionally and molecularly, clearly implicating X-linked gene dosage as the cause of the male/female disparity. Finally, XX female ESCs are also karyotypically unstable, with XO aneuploid cells being rapidly selected for in culture due to their increased fitness compared with XX cells (Choi et al., 2017b; Yagi et al., 2017; Zvetkova et al., 2005).

Despite, or perhaps because of the uniqueness of female ESCs, they are underrepresented in the literature compared to studies performed on male cells. Given the therapeutic potential of ESCs and iPSCs, it is of paramount importance to remedy this and therefore we created the Xmas ESC system. Through the use of the dual fluorescent X-linked reporter alleles in Xmas ESCs we are able to infer both the karyotype and transcriptional status of the female X chromosomes, being the feature that genetically, molecularly and functionally distinguishes female ESCs from males. Moreover, we show through transcriptomic and teratoma experiments, that Xmas ESCs retain normal pluripotency and therefore are an appropriate tool for the study of female-specific pluripotency. Other studies have recently sought to improve the culture of female ESCs, achieving significant stabilisation of the epigenome, including at imprinting control centres, however these culture conditions were unable to preserve the XX karyotype (Choi et al., 2017a; Choi et al., 2017b; Yagi et al., 2017). By producing our X-linked reporter alleles in mice rather than by targeting in ESCs, we were able to ensure a constant supply of *bona fide* XX Xmas ESC lines, rather than cell culture adapted ESCs, enabling us to modify existing methods to develop a protocol that best preserves XX cells. Despite significant improvements however, we were unable to completely stop XO cells arising and becoming predominant in cultured female ESC. Given that the XX karyotype is stable following XCI, the unstable karyotype is likely due to the double dosage of X-linked genes. While we were preparing this manuscript, it was reported that a lower concentration of Mek inhibitor was able to partially stabilise the XX karyotype (Di Stefano et al., 2018). We have tested this and find that low MEK inhibitor further stabilises Xmas ESC karyotype when using our improved conditions (data not shown), but still does not solve the problem entirely. We suggest it may be possible to identify a genetic solution to this problem by performing unbiased screens for genes whose depletion further stabilises the XX karyotype, utilising the Xmas ESC system to efficiently monitor karyotype.

Pluripotency can also be studied *in vitro* through the use of iPSCs and it is these cells that are the hope for the future of regenerative medicine. Interestingly, there appears to be no sex disparity in efficiency of iPSC generation (Di et al., 2015; Kim et al., 2015b), which is perhaps not surprising given that activity from two X chromosomes drives cells towards pluripotency. In culture however, female iPSCs display similar characteristics to ESCs in that they are transcriptionally similar, globally hypomethylated and delayed in their exit from pluripotency compared to XY males. Again, these differences are absent in female XO cells (Pasque et al., 2018; Song et al., 2019). There is therefore a need to also understand female iPSCs as distinct from males, and female Xmas MEFs provide a useful tool for this purpose. Reactivation of the Xi occurs late in the ontogeny of reprogramming and is an indicator of a successfully reprogrammed cell (Pasque et al., 2014). We show here that the Xmas alleles detect reactivation of the Xi at the late stages of reprogramming, therefore Xmas MEFs provide a tractable system to study female-specific reprogramming. In culture female iPSCs with an XO karyotype are rapidly selected, and indeed, in our hands, this occurs even more rapidly in iPSCs than ESCs, likely due to the stresses of the reprogramming process. For female iPSCs to be applied to regenerative medicine, reprogramming and maintenance methods must be optimised to preserve the XX karyotype. A Xmas reporter system made in human cells could expedite this process.

The Xmas ESC system allowed us to culture high quality XX female pluripotent cells for more extended periods of time. We chose to use this system to screen for genes that regulate the establishment of XCI during normal female ESC differentiation. Due to the major issues maintaining XX ESCs, all previous screens for XCI regulators have been performed either in differentiated cells for factors that alter maintenance of XCI (Bhatnagar et al., 2014; Chan et al., 2011; Keniry et al., 2016; Lessing et al., 2016; Li et al., 2018; Minajigi et al., 2015; Minkovsky et al., 2015; Minkovsky et al., 2014; Sripathy et al., 2017), or using non-native (though cunning) systems that instead induce *Xist* out of context, where *Xist* is often in a different chromosomal location in male cells, or in female cells but not induced during the exit from pluripotency (Chu et al., 2015; McHugh et al., 2015; Moindrot et al., 2015; Monfort et al., 2015). Our Xmas ESC system has enabled us to overcome the challenges of working with female ESCs and perform the first screen for regulators of the establishment of XCI in its near-native context. Our screen revealed a role for Smarcc1 and Smarca4 in the establishment of XCI. Both Smarcc1 (also known as Baf155) and Smarca4 (also known as Brg1) are members of the chromatin remodelling BAF (SWI/SNF) complex, with Smarcc1 being the core subunit around which the complex forms (Mashtalir et al., 2018) and Smarca4 being one of a variable number of catalytic ATPase subunits (Wang et al., 1996a; Wang et al., 1996b). Interestingly, the BAF complex is made up of different subunits dependent on cell type, with both Smarcc1 and Smarca4 being part of a ESC-specific complex (known as esBAF), required for both pluripotency and self-renewal (Fazzio et al., 2008; Ho et al., 2009; Kaeser et al., 2008). Our knockdown strategy, designed to avoid disruption of pluripotency by depleting during differentiation, now reveals a new role for esBAF in the exit from pluripotency in females, with both *Smarcc1* and *Smarca4* depletion causing chromosome-wide failure of silencing. Deletion of *Smarcc1* and *Smarca4* in mice are each embryonic lethal peri-implantation, and although consistent with failure of XCI, male embryos also fail to survive, precluding any conclusions being drawn about their role in XCI *in vivo* (Bultman et al., 2000; Han et al., 2008; Kidder et al., 2009). Interestingly, both Smarcc1 and Smarca4 have been found to interact with *Xist* in differentiated cells, with Smarca4 found to have a minor role in the maintenance of XCI when the cells are also challenged with chemical inhibitors of DNA methylation and topoisomerase (Jegu et al., 2019; Minajigi et al., 2015). Here we show a more profound role for Smarcc1 and Smarca4 in the establishment of X chromosome silencing, but no evidence for a role in maintenance of XCI, noting however that our assay includes an inhibitor of DNA methylation but not topoisomerase, suggesting Smarca4 dependent maintenance of XCI could be reliant on topoisomerase.

The major failure of XCI we observe following *Smarcc1* depletion inspired us to produce the first profile of nucleosome occupancy during the establishment of XCI. We have discovered that Smarcc1 is required to deplete promotors of nucleosomes on the future Xi at the very early stages of the establishment of XCI. A similar effect is observed on autosomes and can also be found in published NOMe-seq datasets from the equivalent stages of post-implantation embryos (Argelaguet et al., 2019), suggesting that nucleosome depletion at promoters may be a common occurrence during differentiation. Importantly, however, we have functionally linked this opening to gene silencing; cells with depleted *Smarcc1* fail to open promotors and fail to establish XCI, with the resulting Xi following a similar trajectory to that of the Xa, both in terms of nucleosome positioning and gene silencing. Our data suggest a model where the esBAF complex is recruited to the future Xi following *Xist* upregulation in order to make the Xi accessible to the epigenetic silencing factors required to set up gene silencing, with the complex subsequently excluded from the X once XCI is complete (Figure 7H). Therefore, Smarcc1 sets up a chromatin state necessary for the establishment of silencing. Interestingly, a previous study showed *Xist* both interacted with and repelled Smarca4 in a potentially step-wise fashion (Jegu et al., 2019), suggesting the recruitment and exclusion of Smarca4 from the Xi that we observe may be *Xist* dependent. Broadly, our work revealed a role for nucleosome depletion at the promoter via the esBAF complex in the earliest stages of the XCI, suggesting nucleosome depletion is required to establish silencing. In the future it will be interesting to test what role nucleosome depletion at promoters plays in the initial stages of silencing.

In summary, our new Xmas ESC system has enabled us to optimise the culture of female pluripotent cells, which in turn has allowed us to reveal new requirements for the establishment of XCI. The Xmas cell system provides a renewable resource of high-quality female ESCs and a protocol for optimised culture of such cells that makes the study of female-specific features of pluripotency and differentiation more feasible than ever before.

## Methods

### Key resources table

A list of key resources is provided in Table S7.

### Animal strains and husbandry

Animals were housed and treated according to Walter and Eliza Hall Institute (WEHI) Animal Ethics Committee approved protocols (2014.034, 2018.004). Xmas mice are C57BL/6 background and were maintained as homozygous lines. D4/XEGFP mice were obtained from Jackson labs and backcrossed onto the C57BL/6 background. Xist∆A mice (Royce-Tolland et al., 2010) were obtained from Dr Graham Kay, Queensland Institute of Medical Research, and kept on a 129 background. Castaneus mice were obtained from Jackson labs and maintained at WEHI. FVB/NJ mice were obtained from stocks held at WEHI. Oligonucleotides used for genotyping are provided in Table S8.

### Creation of *Hprt* knockin alleles

The *Hprt* targeted alleles were generated by recombination in Bruce4 C57BL/6 ESCs. The targeting construct was produced by recombineering. This construct was designed to introduce an IRES-mCherry-polyA site or an IRES-eGFP-polyA site sequence 20 bp into the 3’ untranslated region (UTR) of *Hprt*, followed by a PGK-neomycin selection cassette flanked by Frt sites. Note, the mCherry used in the construct contained a synonymous mutation to remove the internal *Nco*I site. The targeting construct also introduced specific sites useful for the Southern blotting strategy used to validate recombination in targeted ESC clones. These sites were *Sph*I and *EcoR*V at the 5’ end, after 20 bp of the 3’ UTR before the IRES, and *EcoR*V and *Nsi*I at the 3’ end before the remainder of the 3’UTR.

Neomycin resistant clones were screened by Southern blot for their 5’ and 3’ integration sites using PCR amplified probes. The 5’ probe was amplified with the 5’-AAACACACACACACTCCACAAA-3’ and 5’-GCACCCATTATGCCCTAGATT-3’ oligos, the 3’ probe was amplified with 5’-GCTGCCTAAGAATGTGTTGCT-3’ and 5’-AAGCCTGGTTTTGGTAGCAG-3’ oligos. Each was cloned into the TopoTA vector. For the Southern blot, DNA was digested individually with *EcoR*V and *Sph*I. The wild-type allele generated a 17.4 kb band with *EcoR*V digestion and the 5’ or 3’ probe, and a 9.2 kb and 8.3 kb knockin band for the 5’ and 3’ probe respectively. The wild-type allele generated a 7.6 kb probe with *Sph*I digestion and the 5’ probe, compared with a 6.4 kb knockin band. The wild-type allele generated an 8.2 kb band with *Nsi*I digestion and the 3’ probe, compared with a 6.7 kb knockin allele.

One Hprt-IRES-mCherry-pA-Frt-neo-Frt and one Hprt-IRES-eGFP-pA-Frt-neo-Frt correctly targeted clone was selected and used for blastocyst injection. The PGK-neo selection cassette was subsequently removed by crossing to the Rosa26-Flpe deleter strain (Farley et al., 2000). The Hprt-IRES-mCherry and Hprt-IRES-GFP alleles were homozygozed and maintained on a pure C57BL/6 background. Genotyping of mice was performed by PCR reaction using GoTaq Green Mix (Promega) and 0.5 μM of each primer, as given in Table S8.

### Derivation and culture of ESCs

Female mice were super-ovulated by injecting 5 IU folligon (MSD Animal Health Australia) two days prior, and 5 IU chorulon (MSD Animal Health Australia) on the day of mating with a stud of the opposite genotype. At E3.5, dams were sacrificed, uteri removed and blastocysts flushed from the uterine horns with M2 medium (Sigma-Aldrich). Blastocysts were washed in M2 medium twice, and 2i+LIF medium [KnockOut DMEM (Life Technologies), 1x Glutamax (Life Technologies), 1x MEM Non-Essential Amino Acids (Life Technologies), 1 X N2 Supplement (Life Technologies), 1 X B27 Supplement (Life Technologies), 1x Beta-mercaptoethanol (Life Technologies), 100 U/mL Penicillin/100 μg/mL Streptomycin (Life Technologies), 10 μg/mL Piperacillin (Sigma-Aldrich), 10 μg/mL Ciprofloxacin (Sigma-Aldrich), 25 μg/mL Fluconazol (Selleckchem), 1000 U/mL ESGRO Leukemia Inhibitory Factor (Merck), 1 μM StemMACS PD0325901 (Miltenyi Biotech), 3 μM StemMACS CHIR99021 (Mitenyi Biotech)] twice. Blastocysts were plated in non-tissue culture treated 24-well plates in 2i+LIF medium. Following 7 days in culture at 37°C in a humidified atmosphere with 5% (v/v) carbon dioxide and 5% (v/v) oxygen, outgrowths were moved by mouth-pipetting through trypsin-EDTA for 2 minutes, ESC wash media [KnockOut DMEM (Life Technologies), 10% KnockOut Serum Replacement (Life Technologies), 100 IU/mL penicillin/100 μg/mL streptomycin (Life Technologies)], and finally 2i+LIF. Outgrowths were disrupted by pipetting and transferred into a 24-well plate to be cultured as ESC lines.

ESCs were maintained in suspension culture in 2i+LIF medium on non-tissue culture treated plates at 37°C in a humidified atmosphere with 5% (v/v) carbon dioxide and 5% (v/v) oxygen. ESCs were passaged daily by collecting colonies and allowing them to settle in a tube for < 5 minutes. Supernatant containing cellular debris was removed and ESC colonies were resuspended in Accutase (Sigma-Aldrich) and incubated at 37°C for 5 minutes to achieve a single-cell suspension. At least 4 x volumes of ESC wash media were added to the suspension and cells were pelleted by centrifugation at 600 x g for 5 minutes, before plating in an appropriately sized non-tissue culture treated plate in 2i+LIF media. Cells were assessed for XX karyotype regularly by flow cytometry.

### Differentiation of ESCs

At least 2 days prior to inducing differentiation ESCs in suspension were allowed to attach by plating onto tissue culture treated plates coated with 0.1% gelatin. Differentiation was induced by transitioning cells from 2i+LIF media into DME HiHi media [DMEM, 500 mg/L glucose, 4 mM L-glutamine, 110 mg/L sodium pyruvate, 15% fetal bovine serum, 100 U/mL penicillin, 100 μg/mL streptomycin, 0.1 mM nonessential amino acids, 50 μM β-mercaptoethanol, and 1000 U/mL ESGRO Leukemia Inhibitory Factor (Merck)] in 25% increments every 24 hours. During this time cells were passaged as required. On the day of transferring into 100% DME HiHi, approximately 10^4^ cells per cm^2^ were plated onto tissue culture treated plates coated with 0.1% gelatin. Cells were not passaged for the remainder of an experiment and media was changed as required.

### Transduction of ESCs

Retrovirus was produced as described (Jansz et al., 2018a; Majewski et al., 2008) and concentrated by precipitation with 4% PEG 8000 followed by centrifugation. ESCs were either seeded at 10^5^ cells per cm^2^ on plates that had been coated with 0.1% gelatin, or at approximately 10^5^ cells per mL in suspension in 2i+LIF medium containing PEG concentrated viral supernatant and 8 μg/mL polybrene. The next day medium was changed, and cells were selected with 1 μg/mL puromycin. shRNA sequences are given in Table S8.

### Teratoma formation

Xmas ESCs were pelleted and washed with PBS before passing through a 70 μm cell strainer. 10^5^ cells were resuspended in 200 μl of 50% Matrigel (Corning) in PBS and injected sub-cutaneous into either the left or right flank of CBA/nude mice. Teratomas were harvested after approximately 60 days, fixed with formalin, embedded in paraffin and stained with Haemotoxylin and Eosin.

### Derivation and culture of MEFs

MEFs were derived from E13.5 embryos and cultured in DMEM supplemented with 10% (v/v) fetal bovine serum at 37°C in a humidified atmosphere with 5% (v/v) carbon dioxide and 5% (v/v) oxygen.

### qRT-PCR

Knockdown efficiency of shRNA retroviral constructs was determined using Roche Universal Probe Library (UPL) assays. Relative mRNA expression levels were determined using the 2^−ddCt^ method, with *Hmbs* as a house-keeping control. Probe numbers and oligonucleotide sequences are provided in Table S8.

### FACS analysis and sorting

Cells were prepared in KDS-BSS with 2% (v/v) FBS and a cell viability dye, 16 μg/mL FluoroGold and analysed using a BD LSRFortesssa cell analyser. Cells were prepared similarly for sorting using a FACSAria. Flow cytometry data were analysed using FlowJo.

Hematopoietic stem and progenitor cells (LSK: Lineage^−^ Sca1^+^ c-Kit^+^ cells) were isolated from fetal livers from E14.5 Xmas female embryos, essentially as described (Kinkel et al., 2015). Dissociated fetal liver cells were incubated with rat monoclonal anti-Ter119 antibody, then mixed with BioMag goat-rat IgG beads (Qiagen) and Ter119^+^ cells were depleted using a Dynal magnet (Invitrogen). The remaining cells were stained with Alexa700-conjugated antibodies against lineage markers Ter119, B220, CD19, Gr1, CD2, CD3 and CD8, APC-conjugated anti-c-kit/CD117 (generated by the WEHI Antibody Facility) and PE-Cy7-conjugated anti-Sca1 (BD Pharmingen). Cells were stained with FluoroGold to assess viability and analysed on a BD LSRFortessa cell analyser.

### X reactivation assay

Xmas or Xi^GFP^Xa^Δ*Xist*^ MEFs were transduced with shRNA retroviruses, selected with 3–5 μg/mL puromycin, then treated with 10 μM 5-azacytidine 3 days post transduction. Cells were analysed by FACS 7 days post transduction. This assay was run exactly as previously described (Keniry et al., 2016).

### iPSC generation

Xmas MEFs were cultured and maintained as previously described (Nefzger et al., 2014). Two days before reprogramming, MEFs were dissociated with 0.25% Trypsin-EDTA (Gibco, 25200114) and labelled (Nefzger et al., 2014) with anti-mouse BUV395 Thy1.2 (BD Biosciences, 565257; 1:200), anti-mouse BV421 EpCAM (BD Biosciences, 563214; 1:100) anti-mouse, SSEA1-Biotin (eBioscience, 13-8813-82; 1:400), Streptavidin Pe-Cy7 (BD Biosciences, 557598; 1:200) and DRAQ7 viability dye (Biolegend, 424001). Using a BD Influx cell sorter (BD Biosciences) setup, GFP^+^/mCherry^−^/Thy1^+^/SSEA-1^−^/EpCAM^−^ cells and GFP^−^/mCherry^+^/Thy1^+^/SSEA^−^1^−^/EpCAM^−^ cells were isolated and seeded onto 0.1% gelatin-coated 6-well plates at 2 x 10^3^ cells per cm^2^. On day −1, Doxycycline-inducible OKSM virus (Millipore, SCR512) and m2rtTA virus (Cyagen Biosciences) were added at a multiplicity of infection of two to cells in MEF medium supplemented with 2 μg/μL Polybrene (Millipore, TR-1003-G). Plates were immediately centrifuged at 750 x g for 60 minutes at room temperature and then incubated at 37°C and 5% CO_2_. At day 0, medium was removed and supplemented with mouse iPSC medium (Nefzger et al., 2014) containing 2 μg/mL Doxycycline (DOX) (Sigma-Aldrich, D9891). Medium was changed every 2 days for 12 days. After day 12 of reprogramming, DOX was withdrawn from culture medium. Cultures were subsequently maintained and passaged regularly with mouse iPSC medium. Cells from reprogramming were harvested on days 3, 6, 9, 12 during reprogramming and iPSC passage 1 (day 16+) for flow cytometry analysis. These cells were labelled with anti-mouse BUV395 Thy1.2 (BD Biosciences, 565257; 1:200), anti-mouse BV421 EpCAM (BD Biosciences, 563214; 1:100) anti-mouse, SSEA1-Biotin (eBioscience, 13-8813-82; 1:400), Streptavidin Pe-Cy7 (BD Biosciences, 557598; 1:200) and DRAQ7 viability dye (Biolegend, 424001). Samples were then analyzed by flow cytometry (Nefzger et al., 2016). For each time point we quantified the percentage of GFP and mCherry positive cells in the populations that were actively undergoing reprogramming by gating in on the time points’ respective reprogramming intermediates as defined in (Nefzger et al., 2014).

### RNA-seq library generation and analysis

For the RNA-seq depicted in Figure 2G,F, Xmas ESCs were derived and cultured as described above and compared to published datasets (Marks et al., 2012; Maza et al., 2015). For the RNA-seq depicted in Figure 3A-C, we derived male C57/Bl6 ESCs using our culture methods, for two independent lines. These cells were then split in two (p0) and cultured for 10 and 20 passages using either the conditions given in this manuscript or the previous state-of-the-art method, described in (Mulas et al., 2019). For the RNA-seq depicted in Figure 5B, Xmas ESC lines were derived and differentiated using the methods described here, with samples collected daily for 8 days of differentiation and compared to published datasets (Marks et al., 2012; Maza et al., 2015). For all *Smarcc1* and *Smarca4* knockdown RNA-seq in female ESCs (Figure 6), we derived female ESCs by crossing FVB/NJ (FVB) dams with CAST/EiJ (CAST) sires. The resultant female ESC lines were expanded and then differentiated using our culture conditions. Cells were transduced with the indicated shRNAs at day 2 of differentiation and samples taken for RNA-seq at the indicated timepoints. For *Smarcc1* and *Smarca4* knockdown RNA-seq in male ESCs (Figure 6J), we derived male C57/Bl6 ESCs and expanded and then differentiated them using our culture conditions. Again, cells were transduced with the indicated shRNAs at day 2 of differentiation and samples taken for RNA-seq at the indicated timepoints.

For all RNA-seq experiments, cells were harvested from plates by the addition of lysis buffer and RNA extracted with a Quick-RNA MiniPrep kit (Zymo Research). Sequencing libraries were prepared using the TruSeq RNA sample preparation kit (Illumina) and sequenced in-house on the Illumina NextSeq500 platform with 75bp reads. For non-allele specific RNA-seq (C57/Bl6 samples), single-end sequencing was performed. Quality control and adapter trimming were performed with fastqc and trim_galore (Krueger) respectively. Reads were aligned to the mm10 reference genome using either tophat (Trapnell et al., 2009) or histat2 (Kim et al., 2015a). Expression values in reads per million (RPM) were determined using the Seqmonk package (www.bioinformatics.babraham.ac.uk/projects/seqmonk/), using the RNA-seq Quantitation Pipeline. Further data interrogation was performed using Seqmonk.

For allele specific RNA-seq (FVBxCAST samples), paired-end sequencing was performed to improve haplotyping efficiency. Quality control and adapter trimming were performed with fastqc and trim_galore (Krueger) respectively. Reads were aligned to a version of mm10 with SNPs between FVB/NJ with CAST/EiJ n-masked, created using SNPsplit (Krueger and Andrews, 2016), using either tophat (Trapnell et al., 2009) or histat2 (Kim et al., 2015a). Reads were haplotype phased using SNPsplit (Krueger and Andrews, 2016) and expression values in reads per million (RPM) determined using the Seqmonk package (www.bioinformatics.babraham.ac.uk/projects/seqmonk/), using the RNA-seq Quantitation Pipeline. For X-chromosome specific analysis, genes were determined to be informative when they had at least 50 mapped and haplotyped reads. Further data interrogation was performed using Seqmonk.

Gene set testing and differential gene expression analysis of male ESC was performed by making two groups by pooling samples at all passages from either the traditional culture method or our improved method. Differential expression analysis between the two ESC culture methods was performed on gene-level counts with TMM normalisation, filtering out genes expressed in fewer than half of the samples, using edgeR v3.26.7 (McCarthy et al., 2012; Robinson et al., 2010). Model-fitting was performed with voom v3.40.6 (Law et al., 2014) and linear modelling followed by empirical Bayes moderation using default settings. Differential expression results from voom were used for gene set testing with EGSEA v1.12.0 (Alhamdoosh et al., 2017) against the c5 Gene Ontology annotation retrieved from MSigDB, aggregating the results of all base methods but ‘fry’ and sorting by median rank.

Distance matrices of differentiating ESCs were determined between gene expression profiles of either *Smarca4* or *Smarcc1* knockdown and the Nons control by calculating the Euclidean distance between log_2_ rpms with the dist function in R v3.6.1

### DNA-seq library preparation and analysis

We derived male C57/Bl6 ESCs using our culture methods, for two independent lines. These cells were split in two (p0) and cultured for 10 and 20 passages using either the conditions given in this manuscript or the previous state-of-the-art method, described in (Mulas et al., 2019). Sequencing libraries were prepared using the TruSeq DNA sample preparation kit (Illumina) and sequenced in-house on the Illumina NextSeq500 platform with 75bp single-end reads. Reads were mapped to mm10 with bowtie2 (Langmead and Salzberg, 2012) and counted in 1Mb bins along the genome using the GenomicAlignments R/Bioconductor package (Lawrence et al., 2013) and computed the percentage of reads mapped to each chromosome. Only bins on the autosomes and sex chromosomes were included and those bins overlapping the ENCODE blacklisted regions were excluded. For each sample, we computed the coverage of each bin in log counts per million. We then computed the log fold changes comparing each sample to the relevant p0 sample and plotted these by bin position along the genome. We used the edgeR R/Bioconductor package (Robinson et al., 2010) to perform a multidimensional scaling plot of distances between samples based on the log fold changes. Differential abundance analysis was performed using edgeR and limma (Ritchie et al., 2015). Briefly, the voom method (Law et al., 2014) was used to prepare count data for linear modelling and the within-cell line correlation estimated using the ‘duplicateCorrelation’ function from the limma package (Smyth et al., 2005). The voom method was then re-applied (now accounting for the within-cell-line correlation), the within-cell-line correlation re-estimated, and these transformed data used as input to a linear model with design matrix encoding the passage number and protocol of each sample while blocking on the cell line and including the estimated within-cell-line correlation when fitting the linear models. We used the empirical Bayes statistics (Phipson et al., 2016) to test for differential abundance at p10 vs. p0 and p20 vs. p0 within each protocol at a false discovery rate of 0.05 and requiring a minimum log2-fold change of 1.1 (McCarthy and Smyth, 2009).

### Immunofluorescence

Immunofluorescence was performed as described in (Chaumeil et al., 2008), with modifications on differentiating C57/Bl6 female ESCs at day 6. Cells were fixed with 3% (w/v) paraformaldehyde in PBS for 10 min at room temperature, washed 3 times in PBS for 5 minutes each and permeabilised in 0.5% (v/v) triton X-100 for 5 minutes. Cells were blocked in 1% (w/v) Bovine serum albumin (BSA) in PBS for 20 minutes, then incubated in primary antibody in the 1% (w/v) BSA overnight at 4°C in a humid chamber. Primary antibodies used were Smarca4 (1:100 ab110641, Abcam) and H3K27me3 (1:100 07-449, Millipore or 1:100 C36B11, Cell Signalling Technology). Cells were washed three times in PBS for 5 minutes each and then incubated with a secondary antibody diluted in 1% (w/v) BSA for 40 minutes at room temperature in a dark, humidified chamber. Secondary antibodies used were Donkey anti-rabbit IgG Alexa Fluor 555 conjugate (1:500, A315Thermo Fisher) and Goat anti-rabbit IgG Alexa Fluor 647 conjugate (1:500, A21244 Thermo Fisher). For the simultaneous staining of Smarcc4 and H3K27me3, H3K27me3 (C36B11) rabbit mAb Alexa fluor 647 conjugate (Cell Signalling Technology) was used after the secondary antibody was washed off and incubated for 1 hour in a dark humidified chamber at room temperature. Nucleus was stained with DAPI (0.2 μg/mL) in PBS for 5 minutes at room temperature. Cells were mounted in Vectashield antifade mounting medium (Vector Laboratories) and visualised on LSM 880 or LSM 980 microscopes (Zeiss). Image analysis was performed in a semi-automated fashion using a custom written Fiji (Schindelin et al., 2012) macro. The researcher was presented with an image and manually segmented the cells of interest using the region manager. Auto-thresholding methods were used to segment the nuclei and the H3K27me3 region, and mean intensity of Smarca4 measured in both the whole nucleus and region containing H3K27me3.

### *Xist* RNA fluorescence *in situ* hybridisation (FISH)

*Xist* RNA FISH was performed as previously described (Chaumeil et al., 2008; Jansz et al., 2018b) on day 4 or day 5 in differentiated C57/Bl6 female ESCs. *Xist* RNA was detected with a 15 kb cDNA, *pCMV-Xist-PA*, as previously described (Wutz and Jaenisch, 2000). The *Xist* probe was labelled with Green-dUTP (02N32-050, Abbott) by nick translation (07J00-001, Abbott). The cells were mounted in Vectashield antifade mounting medium (Vector Laboratories) and visualised on LSM 880 or LSM 980 microscopes (Ziess). Images were analysed using the open source software FIJI (Schindelin et al., 2012).

### NOMe-seq library generation and analysis

Female ESCs were derived by crossing FVB/NJ dams with CAST/EiJ sires. The resultant female ESC lines were expanded and then differentiated using our culture conditions. Cells were transduced with the indicated shRNAs at day 2 of differentiation and samples fixed in 1% formaldehyde at the indicated timepoints. NOMe-seq samples were prepared as described (Lay et al., 2018), following their protocol for fixed cells. Bisulfite treatment was performed using the EZ DNA Methylation kit (Zymo Research) and sequencing libraries prepared with the Accel-NGS Methyl-Seq DNA Library Kit (Swift Biosciences) and sequenced in-house on the Illumina NextSeq500 platform with 75bp paired-end reads. Quality control and adapter trimming were performed with fastqc and trim_galore (Krueger) respectively. Using bismark (Krueger and Andrews, 2011), reads were aligned to a version of mm10 with SNPs between FVB/NJ with CAST/EiJ n-masked, created using SNPsplit (Krueger and Andrews, 2016) then bisulfite converted using bismark. Reads were haplotype phased using SNPsplit (Krueger and Andrews, 2016) and methylation calls made with the bismark_methylation_extractor (Krueger and Andrews, 2011). Methylation calls were filtered for informative CpG and GpC positions using coverage2cytosine with the --nome-seq flag. For analysis of GpC methylation, % methylation was determined at all covered GpC positions and then averaged over 25 positions and normalised using Enrichment normalisation with the Seqmonk package (www.bioinformatics.babraham.ac.uk/projects/seqmonk/). Both heatmap and line plots were produced by averaging over all gene positions in the indicated genomic regions, with line graphs additionally smoothed for clarity using Seqmonk.

### Accession Numbers

All next generation sequencing data generated for this project have been deposited in the Gene Expression Omnibus (GEO) database under accession number GSE137163.

## Acknowledgements

This study was supported by an Australian Research Training Program scholarship (NJ, LJG), a Melbourne Research Scholarship - International (IW), the Bellberry-Viertel Senior Medical Research Fellowship (MEB), Sylvia and Charles Viertel Senior Medical Research Fellowship (JMP) and an Australian National Health and Medical Research Council fellowship (MER). Grant support was provided by the Australian National Health and Medical Research Council (1059624 to MEB, 1140976 to MEB, MER and AK), the Dyson Bequest and the DHB Foundation. Additional support was provided by the Victorian State Government Operational Infrastructure Support, Australian National Health and Medical Research Council IRIISS grant (9000433).

## Contributions

AK, LJG, DJH and MEB conceived the study. AK, NJ, LJG, IW, JC, CMN, JL, KAB, MI, TB, ATdF, TW and MP performed experiments. AK, PFH, QG and LW performed bioinformatic analysis. SAK and PCT provided expertise. AK, MER and MEB secured funding. MER, JMP and MEB supervised the project. AK and MEB wrote the manuscript with contribution from all authors.

## Declaration

The authors declare no competing interests.

---

**Figure S1.**
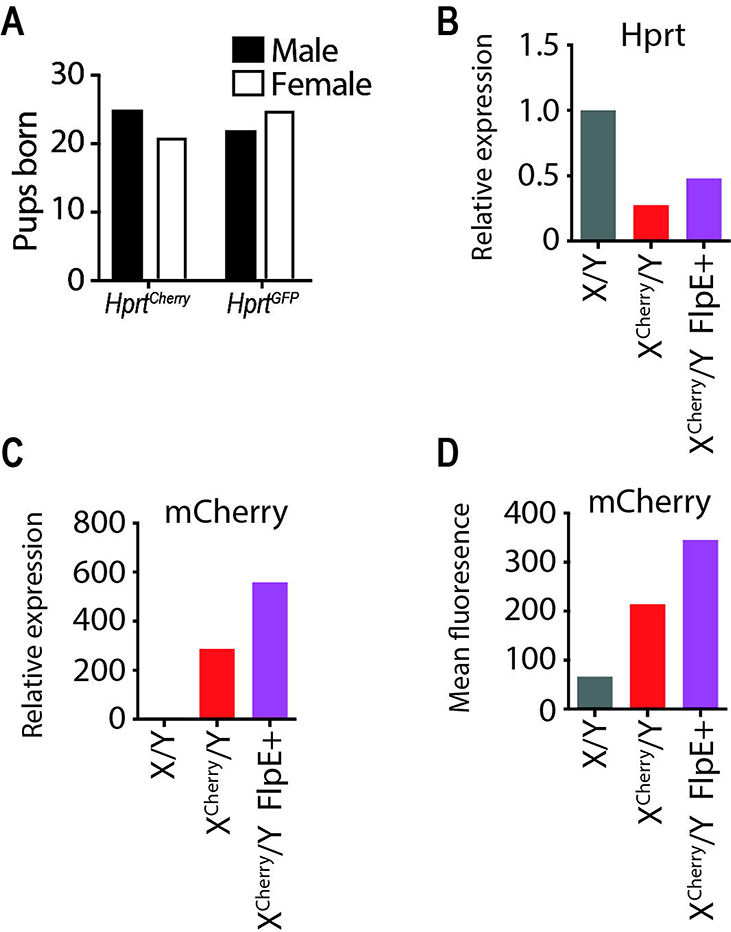
Xmas reporter alleles are not detrimental to survival of female mice. **(A)** Numbers of male and female mice born of the indicated homozygous/hemizygous genotypes. **(B)** *Hprt* expression measured by qRT-PCR in ESCs of the indicated genotypes. **(C)** mCherry expression measured by qRT-PCR in ESCs of the indicated genotypes. **(D)** mCherry expression measured by flow cytometry in ESCs of the indicated genotypes.

**Figure S2.**
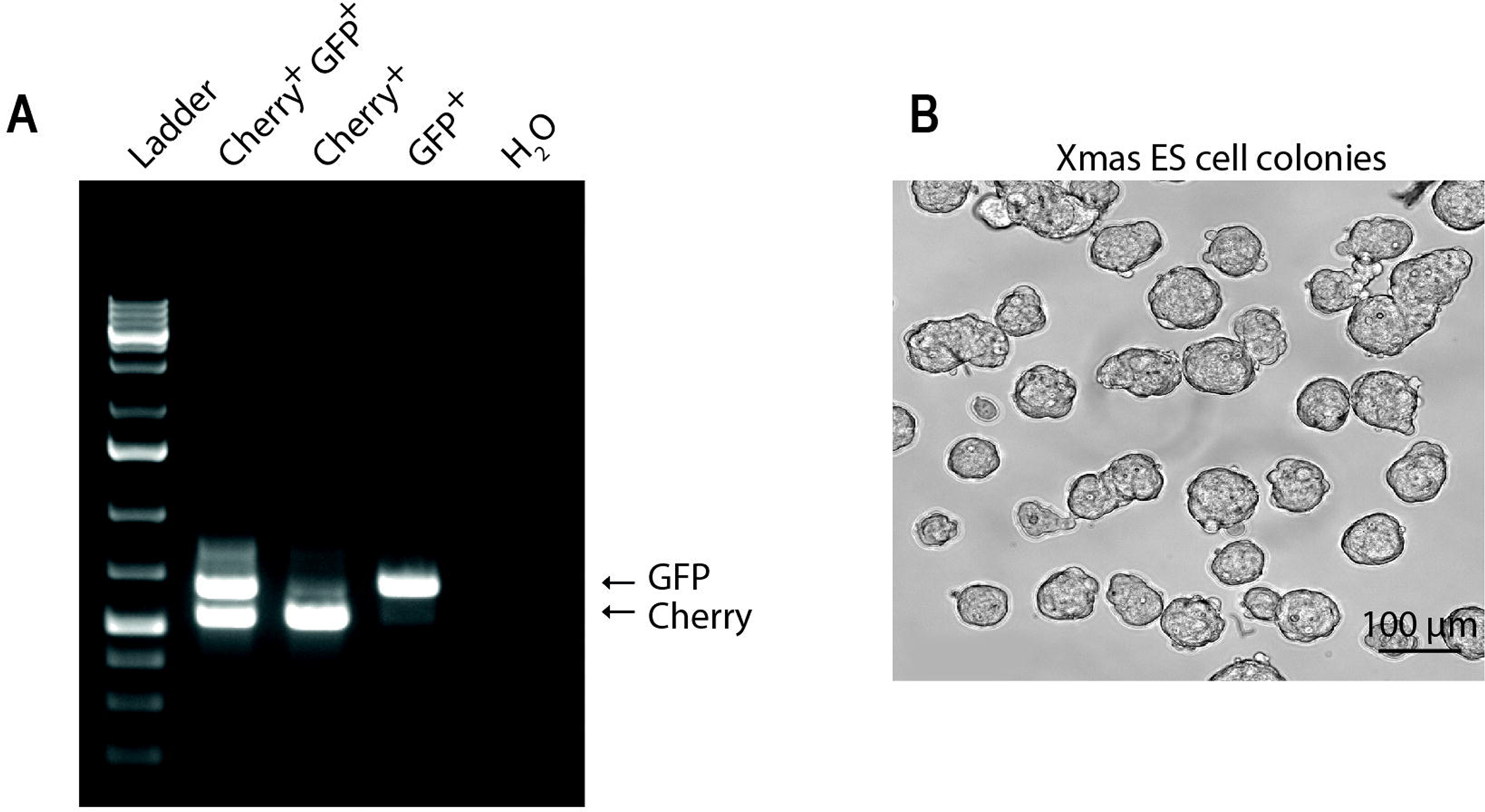
Xmas ESCs allow development of improved culture conditions. **(A)** Gel electrophoresis of PCR product of the fluorescent reporter constructs produced from DNA of Xmas ESCs purified by FACS into GFP^+^, mCherry^+^ and Cherry^+^GFP^+^ double positive populations. **(B)** Bright field microscopy image of Xmas ESC colonies maintained under our improved culture conditions.

**Figure S3.**
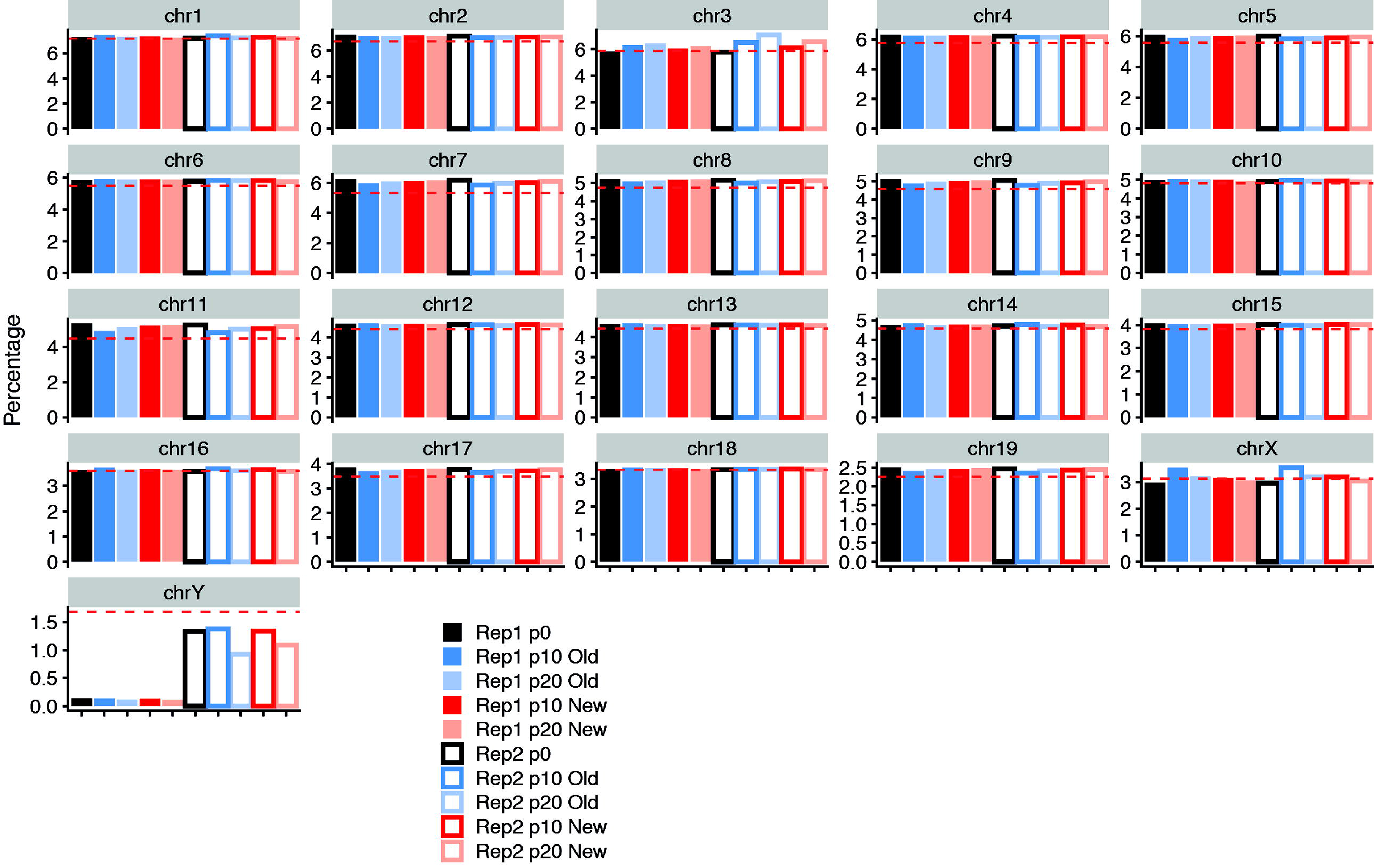
Male ESCs are prone to losing a Y chromosome in culture. Bar graphs showing DNA-seq data from male ESCs maintained in 2i media with the percentage of total reads mapping to the indicated chromosomes at p0 or following p10 or p20 passages in either traditional (Old) or our improved (New) culture conditions. Dotted lines indicate the expected percentage of reads that should map to the chromosome based on chromosome length. Data for two independent male ESC lines is shown.

**Figure S4.**
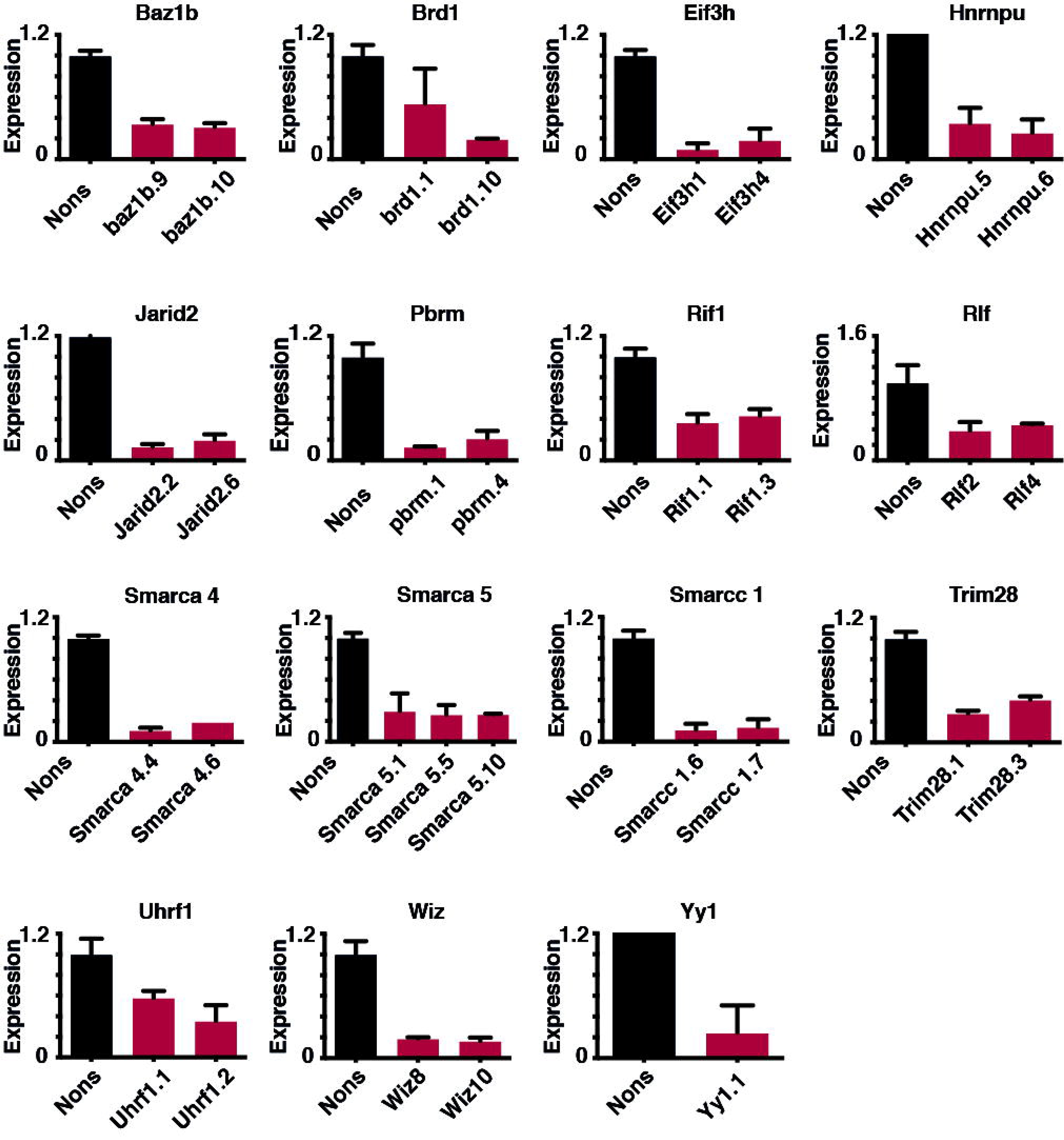
Hairpin validation. Bar graphs showing the expression of the indicated genes relative to *Hmbs*, measured by qRT-PCR following knockdown with the indicated shRNAs. Knockdown was measured in either ESCs or MEFs three days following viral transduction of shRNA (n>3, error bars indicate s.e.m.).

**Figure S5.**
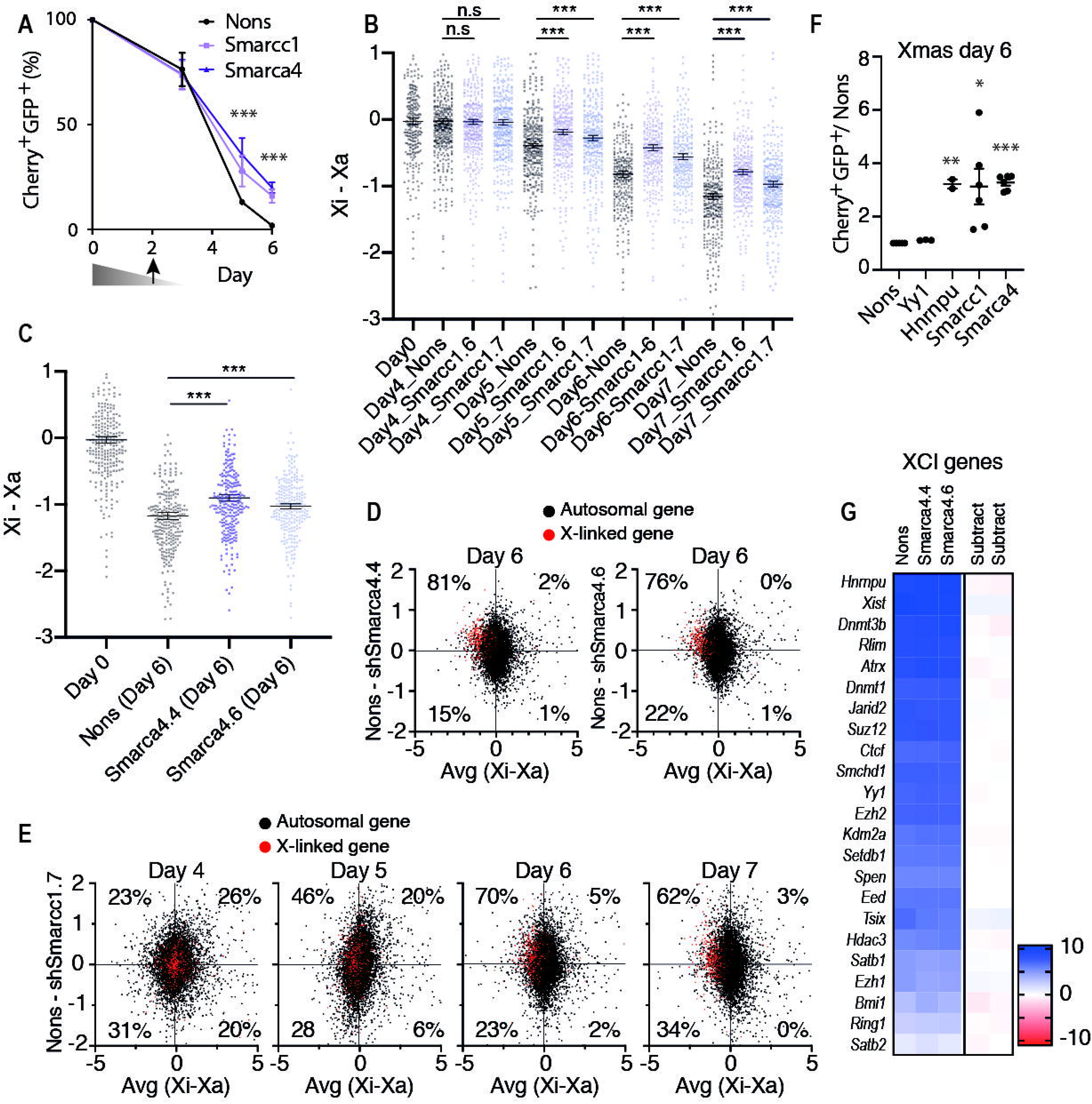
*Smarcc1* and *Smarca4* depletion cause failure of XCI. **(A)** Flow cytometry data along a time course of Xmas ESC differentiation following shRNA mediated knockdown of *Smarcc1*, *Smarca4* or Nons (n = 4 for each of two independent shRNAs per gene, error bars indicate s.e.m., Student’s paired *t*-test, *** indicates *p* < 0.001). **(B,C)** Expanded version of data from Figure 6D,E showing allele specific RNA-seq of differentiating X^FVB^X^CAST^ ESCs following knockdown with indicated hairpins against *Smarcc1* **(B)** and *Smarca4* **(C)**. Each point represents the Xi-Xa log_2_ expression value of an individual informative X-linked gene (error bars indicate s.e.m., Student’s paired *t*-test, *** indicates *p* < 0.001). **(D,E)** These graphs show RNA-seq data and are designed to compare gene expression from the X chromosome with respect to autosomes. Each point on the graph represents an informative gene, with X-linked genes in red and autosomal genes in black. The x-axis shows the ratio of expression from FVB compared to CAST (Xi-Xa log_2_), therefore XCI is observed as a left shift of the red dots along the x-axis. The y-axis shows the ratio of expression from Nons compared to knockdown with *Smarca4* **(D)** or *Smarcc1* **(E)** (log_2_FC rpm Nons − knockdown), therefore a failure of XCI in the knockdown is observed as an upward shift of the red dots along the y-axis. Black dots give an indication of global trends in gene expression. Dotted lines indicate medians and percentages show the X-linked genes falling into each quadrant. **(F)** Xmas ESCs transduced with the indicated hairpins on day 3 of differentiation, with fluorescence measured by flow cytometry at day 6. (n = 2-6 error bars show the s.e.m., Student’s paired *t*-test, *, **, *** indicate *p* < 0.05, *p* < 0.01, *p* < 0.001 respectively. **(G)** Heat map of gene expression (rpm log_2_) of known regulators of XCI, with the difference between knockdown and control (subtract, Nons – knockdown) indicated.

**Figure S6.**
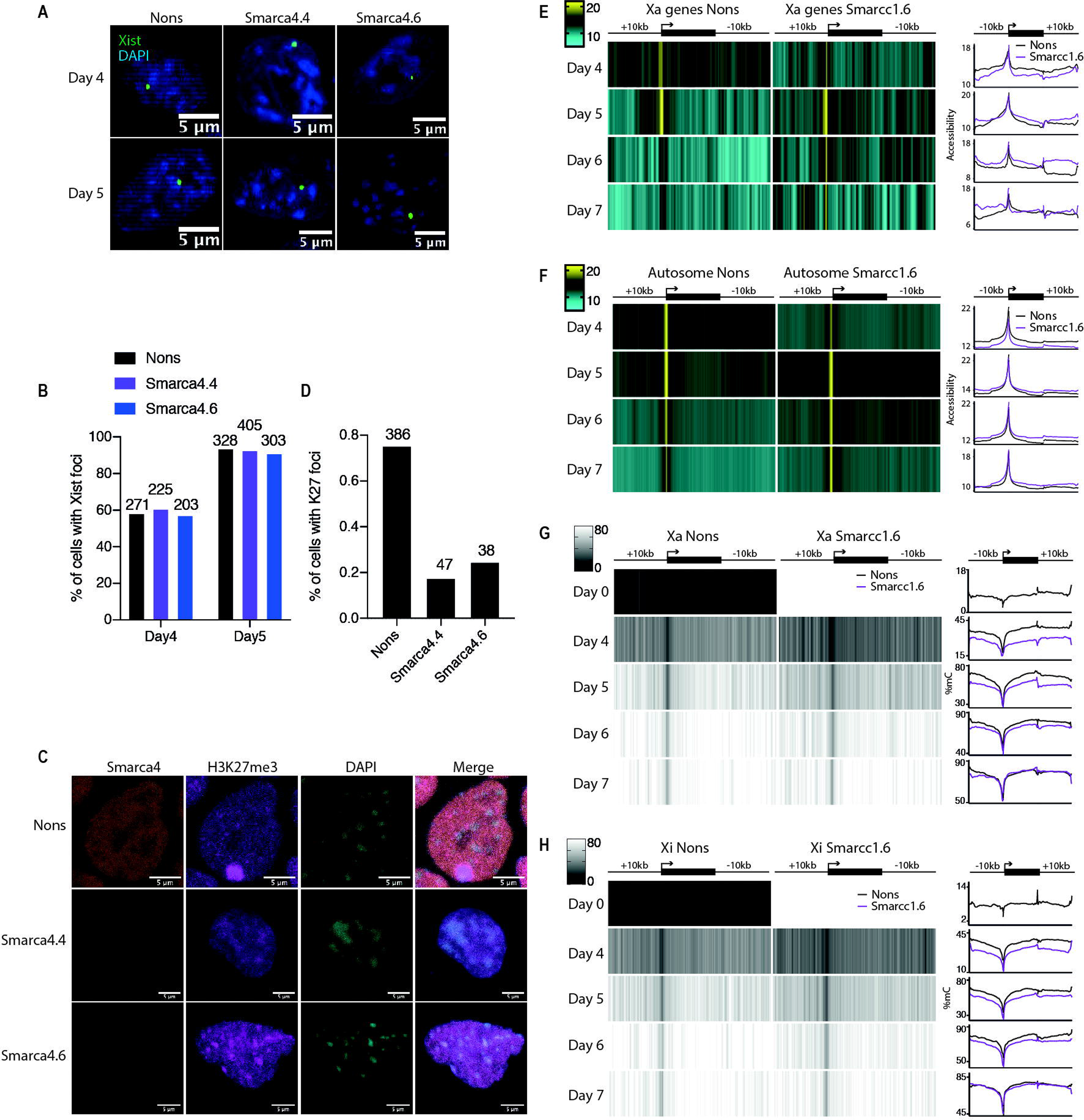
Smarcc1 and Smarca4 are required to establish XCI. **(A)** RNA FISH for *Xist* in female ESCs at day 4 and 5 of differentiation following knockdown with the indicated hairpins. **(B)** Quantification of data from (A). **(C)** Immunostaining of H3K27me3 and Smarca4 in female ESCs at day 6 of differentiation following knockdown with the indicated hairpins. **(D)** Quantification of data from (C). **(E,F)** Nucleosome occupancy (% GpC methylation) along a time course of female ESC differentiation determined by NOMe-seq averaged across all genes and flanking regions on the Xa **(E)** or autosomes **(F)** upon Smarcc1.6 knockdown. The data shown as a heat map or a smoothed histogram. **(G,H)** DNA methylation (% CpG methylation) along a time course of female ESC differentiation determined by NOMe-seq averaged across all genes and flanking regions on the Xa **(E)** or Xi **(F)** upon Smarcc1.6 knockdown. The data shown as a heat map or a smoothed histogram.

**Figure S7.**
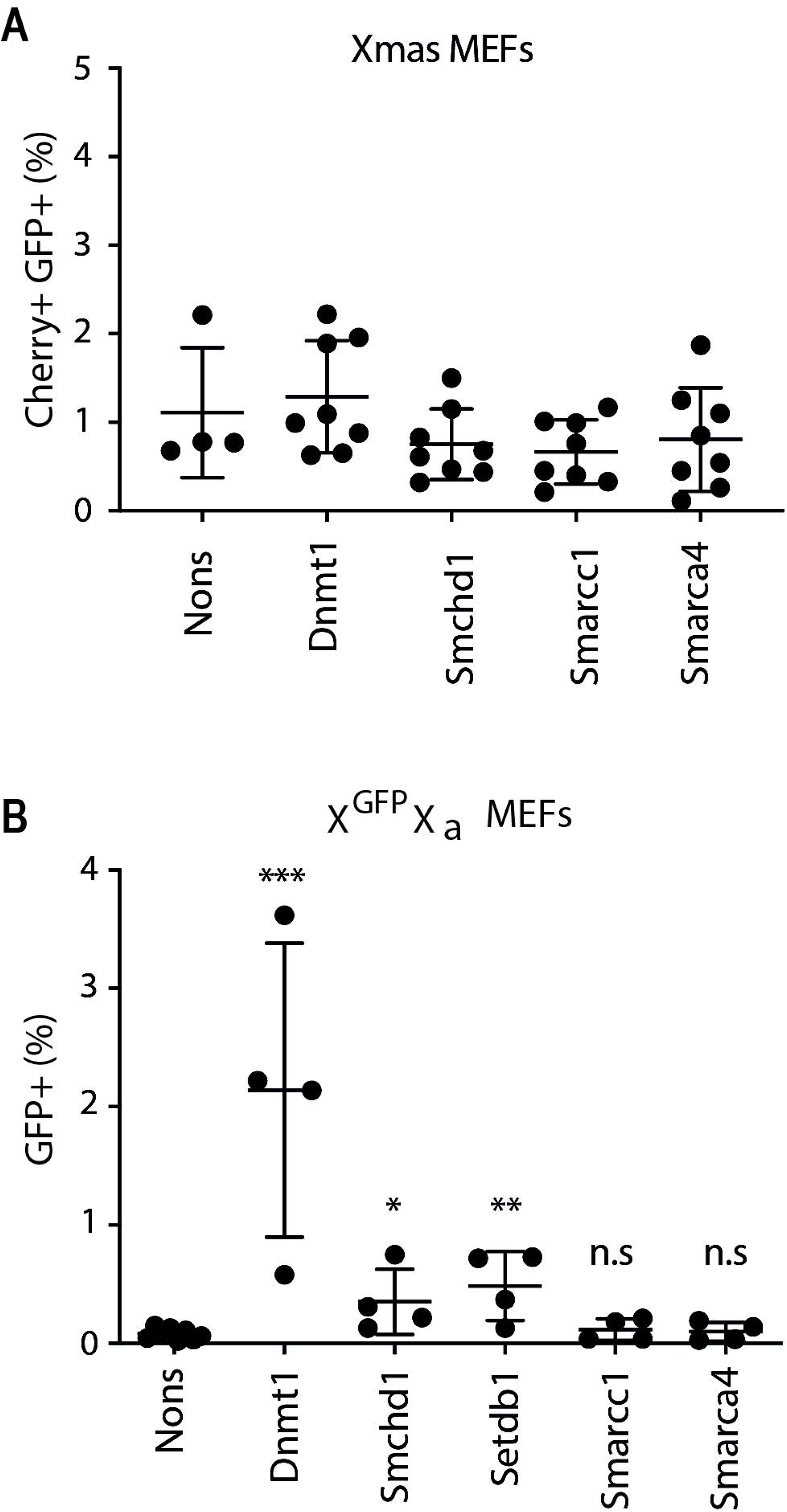
Smarcc1 and Smarca4 are not required for maintenance of XCI. **(A)** Flow cytometry data from Xmas MEFs following knockdown with two independent hairpins per gene and treatment with 5-azacytidine. n = 4–8 error bars show the s.e.m. **(B)** Flow cytometry data from Xi^GFP^Xa MEFs following knockdown with two independent hairpins per gene and treatment with 5-azacytadine. n = 4–8 error bars show the s.e.m., Student’s unpaired *t*-test, *, **, *** indicate *p* < 0.05, *p* < 0.01, *p* < 0.001 respectively.

**Table S1 Differential gene expression analysis of RNA-seq data from male ESC** Table provides the results of differential gene expression between male ESCs cultured with either our improved culture conditions (New) or traditional methods (Old). p10 and p20 samples have been considered replicates in this analysis.

**Table S2 Differential genomic representation analysis of DNA-seq data from male ESC** Table provides the results of differential genomic representation of 1Mb bins between male ESCs cultured with either our improved culture conditions (New) or traditional methods (Old). The logFC value and the adj.P.Val compares the indicated sample with the equivalent p0 sample.

**Table S3 RNA-seq in differentiating Xmas ESCs** Table provides analysed RNA-seq data along a timecourse of Xmas ESC differentiation. Expression values are given in rpm log_2_.

**Table S4 RNA-seq in female ESC with Smarca4 knockdown** Table provides allele specific RNA-seq data at day6 of FVB cross CAST female ESC differentiation with either Smarca4 knockdown or Nons control. Values given are the Xi-Xa log_2_ value for only informative genes.

**Table S5 RNA-seq in female ESC with Smarcc1 knockdown** Table provides allele specific RNA-seq data along a timecourse of FVB cross CAST female ESC differentiation with either Smarcc1 knockdown or Nons control. Values given are the Xi-Xa log_2_ value for only informative genes.

**Table S6 RNA-seq in male ESC** Table provides analysed RNA-seq data along a timecourse of male ESC differentiation. Expression values are given in rpm log_2_.

**Table S7 Key resources** Table provides a list of key resources used in this study.

**Table S8 Oligonucleotides** Table provides the sequences of oligonucleotides used in this study for qRT-PCR, genotyping and shRNA knockdown.

